# Active repression of cell fate plasticity by PROX1 safeguards hepatocyte identity and prevents liver tumourigenesis

**DOI:** 10.1101/2024.09.10.612045

**Authors:** Bryce Lim, Aryan Kamal, Borja Gomez Ramos, Juan M. Adrian Segarra, Ignacio L. Ibarra, Lennart Dignas, Tim Kindinger, Kai Volz, Mohammad Rahbari, Nuh Rahbari, Eric Poisel, Kanela Kafetzopoulou, Lio Böse, Marco Breinig, Danijela Heide, Suchira Gallage, Jose E. Barragan Avila, Hendrik Wiethoff, Ivan Berest, Sarah Schnabellehner, Martin Schneider, Jonas Becker, Dominic Helm, Dirk Grimm, Taija Mäkinen, Darjus F. Tschaharganeh, Mathias Heikenwalder, Judith B. Zaugg, Moritz Mall

## Abstract

Cell fate plasticity enables development, yet unlocked plasticity is a cancer hallmark. Regulating cell identity requires gene activation and repression. While master regulators induce lineage-specific genes to restrict plasticity, it remains unclear whether unwanted plasticity is actively suppressed by lineage-specific repressors. Here, we computationally predict so-called safeguard repressors for 18 cell types that block phenotypic plasticity lifelong. We validated hepatocyte-specific candidates using reprogramming, revealing that Prospero homeobox protein 1 (PROX1) enhanced hepatocyte identity by direct repression of alternate fate master regulators. In mice, Prox1 was required for efficient hepatocyte regeneration after injury and acted as a tumour suppressor in multiple liver cancer models. In line with patient data, *Prox1* depletion caused hepatocyte fate loss *in vivo*, and promoted transition of hepatocellular carcinoma to cholangiocarcinoma, conversely, overexpression promoted cholangiocarcinoma to hepatocellular carcinoma transdifferentiation. Our findings provide mechanistic evidence for PROX1 as a hepatocyte-specific safeguard and support a model where individual cell type-specific repressors actively suppress plasticity throughout life to safeguard lineage choice and prevent disease.

## Main

Cell fate plasticity enables stem and progenitor cells to generate all cell types of the body^1^. During differentiation, this plasticity typically is restricted to reach a terminally-differentiated state with a defined phenotype. Conversely, unlocked cellular plasticity emerged as a cancer hallmark^2^. In this context, blocked differentiation or even de- and trans-differentiation of mature cells to a malignant state contributes to neoplasia. Precise mechanisms that govern phenotypic plasticity often remain elusive, but transcription factors are important regulators of this process.

Individual lineage-specific selectors or master regulators can activate gene regulatory networks to induce specific tissues or cell types^3–6^. Loss of master regulators in mature cells, such as deletion of *Pax5* in B cells or *Ptf1a* in acinar cells^7,8^, can cause cancer. Repressors that silence unwanted genes are also important to define cell fate in development^9,10^. A prominent example is the developmental repressor REST^11^, which represses neuronal genes in non-neuronal cells. However, one repressor silencing one cell identity raises a logistical problem, since hundreds of repressors would be needed to silence all alternate identities in each cell type. We recently discovered a new kind of cell type-specific “safeguard repressor” that might resolve this conundrum. Unlike REST, the transcription factor MYT1L is almost exclusively expressed in neurons and directly binds and represses several non-neuronal genes to promote neuronal cell identity^12,13^. MYT1L is expressed lifelong, and loss of MYT1L is associated with mental disorders and brain cancer^14,15^. Whether this type of repression exists in other lineages, and how it contributes to cellular plasticity in cancer, is unclear.

Here, we developed an *in silico* screen to identify cell type-specific safeguard transcription factors across 18 cell types that promote and maintain cell identity by actively suppressing cell fate plasticity and cancer. We experimentally validated hepatocyte-specific candidates, revealing that PROX1 enhanced hepatocyte identity by repressing alternate cell fate master regulators during reprogramming. In mice, *Prox1* was required for efficient hepatocyte regeneration after injury and sufficient to block liver tumour initiation and progression. Interestingly, manipulating PROX1 levels can toggle transformed hepatocytes between cholangiocarcinoma and hepatocellular carcinoma fates. Together, these findings support a model in which cell type-specific safeguard repressors actively suppress unwanted cell fate plasticity to induce and maintain cell identity and block tumourigenesis.

## Results

### Identifying safeguard repressor candidates across eighteen cell types

To globally identify safeguard repressors that can suppress cell fate plasticity and maintain cell identity by actively silencing alternative fates, we defined three cardinal features they should: i) exhibit lifelong and cell type-specific expression; ii) bind and repress genes expressed in other cell types; and iii) promote and maintain a particular cell identity (**Fig. 1a**). To identify candidates, we focused on 18 well-characterised cell types spanning all three germ layers and used expression data from Tabula Muris, a single-cell gene expression atlas derived from multiple tissues of approximately three-month-old adult mice^16^ (**Extended Data Fig. 1a**). First, we calculated cell type-specific expression for all 1,296 transcription factors detected across the 18 cell types. Next, we created cell type-specific gene expression signatures for all 18 cell types (**Extended Data Fig. 1b; Supplementary Table 1**). For each transcription factor we then counted the number of binding motifs in the promoters of these signature genes. For safeguard repressor candidates we expect that their DNA-binding motifs are depleted at signature genes of the desired cell type and enriched at alternative fate genes. Finally, we integrated transcription factor expression levels and DNA-binding motif depletion at signature genes to derive a safeguard repressor score for each transcription factor in each cell type (**Extended Data Fig. 1c; Methods**).

**Fig. 1:**
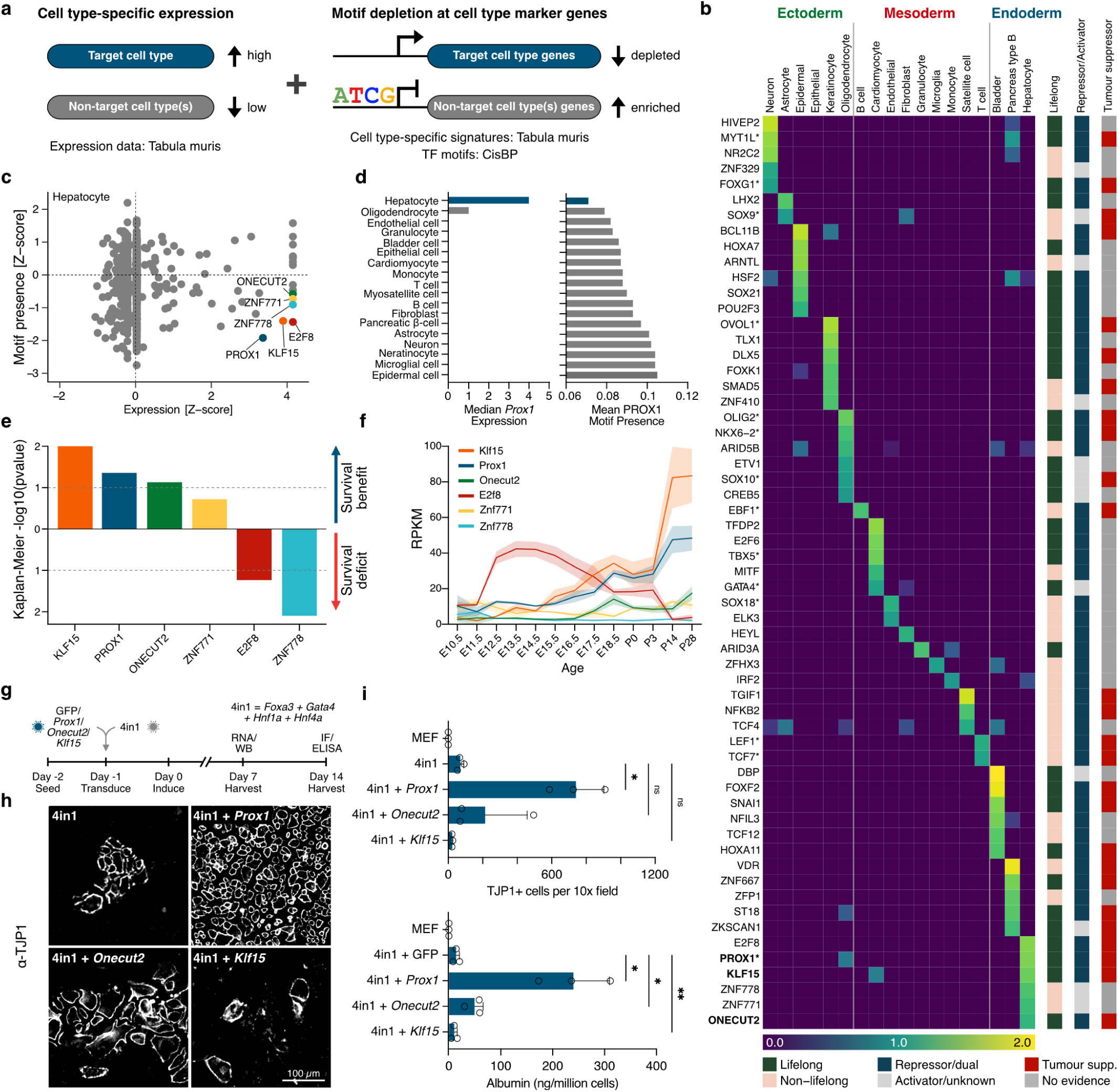
Safeguard repressors revealed by a computational and experimental screen. a, Transcription factors with high expression in the target cell type, and motif depletion in target cell type-specific promoters, have high safeguard repressor scores (see also **Extended Data Fig. 1c)**. b, Top six safeguard repressor candidates across eighteen cell types based on safeguard repressor score > 0. Shown are lifelong expression based on Tabula Muris Senis, repressor/activator activity, and tumour suppressor role from published reports. Asterisks: factors reported to promote indicated cell fate (**Supplementary Table 2**) Bold: candidates in this study. c, Expression and motif presence analysis of 1,296 transcription factors highlights six hepatocyte safeguard repressor candidates. d, Left, *Prox1* expression in 18 cell types from Tabula Muris. Right, number of PROX1 motifs in promoters of cell type marker genes. e, Predicted survival benefit or deficit based on liver candidate expression in tumours. Log rank test based on Kaplan-Meier curves from hepatocellular carcinoma patients in TCGA, segregated by high or low expression of each candidate (see also **Extended Data Fig. 1h**). f, Developmental expression of the top six hepatocyte repressor candidates in mouse liver^18^. g, Experimental validation of safeguard repressor candidates by lentiviral overexpression during 4in1-induced hepatocyte (iHep) reprogramming from mouse embryonic fibroblasts (MEFs). h, Representative TJP1 immunofluorescence images of induced hepatocytes following overexpression of top three hepatocyte repressor candidates or GFP control. i, Number of TJP1+ cells based on immunofluorescence quantification (top) and Albumin secretion from ELISA measurements (bottom) at day 14 of iHep reprogramming with indicated candidates. Bar graphs show mean values from three biological replicates, error bars = SD, Dunnett’s test, * p-adj < 0.05, ** p-adj < 0.01.

Following this approach, we shortlisted 59 candidates across 18 cell types (**Fig. 1b**). 50 of these transcription factors have reported repressor or dual activator/repressor function based on literature evidence (**Supplementary Table 2**). We further classified 33 candidates as lifelong-expressed based on continued expression in approximately two-year-old mice from Tabula Muris Senis^17^. Indeed, using a developmental gene expression atlas for heart, brain, and liver tissues we found that 77% (17/22) of the candidates were expressed at a high level throughout life in the respective cell type^18^, which is a significant enrichment compared to all transcription factors expressed in these organs (p = 6.25e-5) (**Supplementary Table 2**). Across all analysed germ layers and cell types, 27 candidates satisfied our criteria for lifelong safeguard repressors. Based on previous work, 14 of these candidates promote the predicted cell fate during development or reprogramming (**Supplementary Table 2**), and some act, at least in part, as repressors, including TBX5 in cardiomyocytes and OLIG2 in oligodendrocytes^19,20^. Our screen also identified MYT1L as a safeguard repressor, confirming our previous experimental evidence that MYT1L induced and maintained neuronal identity by actively repressing non-neuronal genes (**Fig. 1b**; **Extended Data Figs. 1d-f**)^12,13^. For the top hepatocyte candidate PROX1, we generated chromatin binding data from primary mouse liver using CUT&RUN that confirmed binding enrichment at many non-hepatocyte specific genes corroborating the results from our motif-based screen (**Figs. 1c-d**; **Extended Data Fig. 1g; Supplementary Table 2**). Interestingly, 59% (16/27) of the candidates are reported to have tumour-suppressive roles in their respective cell types, this enrichment was especially prominent for endodermal candidates including in hepatocytes (**Fig. 1b; Supplementary Table 2**).

### PROX1 is the most biologically-relevant hepatocyte-specific safeguard repressor

Cell fate loss plays a crucial role in liver disease, including dedifferentiation and transdifferentiation in the development of primary liver cancer^21,22^. Thus, we analysed the expression of our hepatocyte-specific candidates in hepatocellular carcinoma (HCC) patients from The Cancer Genome Atlas (TCGA). Of the top six predicted safeguard repressors, high expression of *PROX1*, *KLF15, ONECUT2,* and *ZNF771* was associated with better prognosis, suggesting they could function as putative liver tumour suppressors (**Fig. 1e**; **Extended Data Fig. 1h; Methods**). However, only *Prox1*, *Onecut2*, and *Klf15* showed high lifelong expression throughout development and ageing in the liver (**Fig. 1b,f; Supplementary Table 2**).

To test whether our top three candidates promote hepatocyte identity, we studied their effects in the context of cell fate reprogramming. Previous work showed that overexpression of four liver transcription factors – *Foxa3*, *Gata4*, *Hnf1a,* and *Hnf4a* – from a single polycistronic vector (4in1) reprogrammed mouse embryonic fibroblasts (MEFs) towards induced hepatocytes^23^. We thus assessed the effects of *Prox1*, *Onecut2*, or *Klf15* overexpression during hepatocyte reprogramming (**Fig. 1g**). While all candidate factors were expressed at a similar level, only PROX1 increased expression levels of several hepatocyte-specific proteins and genes, such as *Cdh1* and *Krt18* **(Extended Data Figs. 1i-l)**. Indeed, compared to baseline reprogramming, PROX1 overexpression generated >10-fold more hepatocyte-like cells based on morphology and TJP1 expression, and increased Albumin secretion per cell 17-fold (**Figs. 1h,i**). PROX1 is a Prospero homeobox transcription factor that plays a role in the development of several tissues, including the liver^24^. The striking promotion of hepatocyte cell fate by PROX1 during reprogramming and the association of high expression with better survival in HCC patients encouraged us to investigate the role of PROX1 as a potential liver tumour suppressor.

### *PROX1* expression positively correlates with survival in hepatocellular carcinoma patients

The role of PROX1 in liver cancer is contradictory, as some studies indicate a tumour-promoting function, while others propose a tumour suppressor role^25,26^. Thus, we decided to examine the role of PROX1 as a putative hepatocyte cell fate safeguard in liver cancer. We first assessed whether *PROX1* expression is dysregulated in HCC patients^27^. We found that expression in tumour samples (n=62) was lower than in paired normal tissue controls (n=59) (**Fig. 2a**). Immunohistological analysis confirmed that PROX1 protein levels are significantly reduced in HCC patient tumours compared to the adjacent non-tumour liver tissue (**Fig. 2b**; **Extended Data Fig. 2a**). We then analysed whether *PROX1* expression levels in HCC patients correlated with disease prognosis. In a cohort of 364 patients with both transcriptome and survival data^28^, we found that patients with high *PROX1* expression had a median survival of 81.9 months, while patients with low *PROX1* levels had a median survival of only 47.4 months (**Fig. 2c**). Strikingly, we found that chromosomal amplifications that encompass *PROX1* are associated with increased survival in HCC patients^29–35^ (**Fig. 2d)**. These genetic alterations and expression changes suggest that PROX1 has a tumour-suppressive role in HCC patients.

**Fig. 2:**
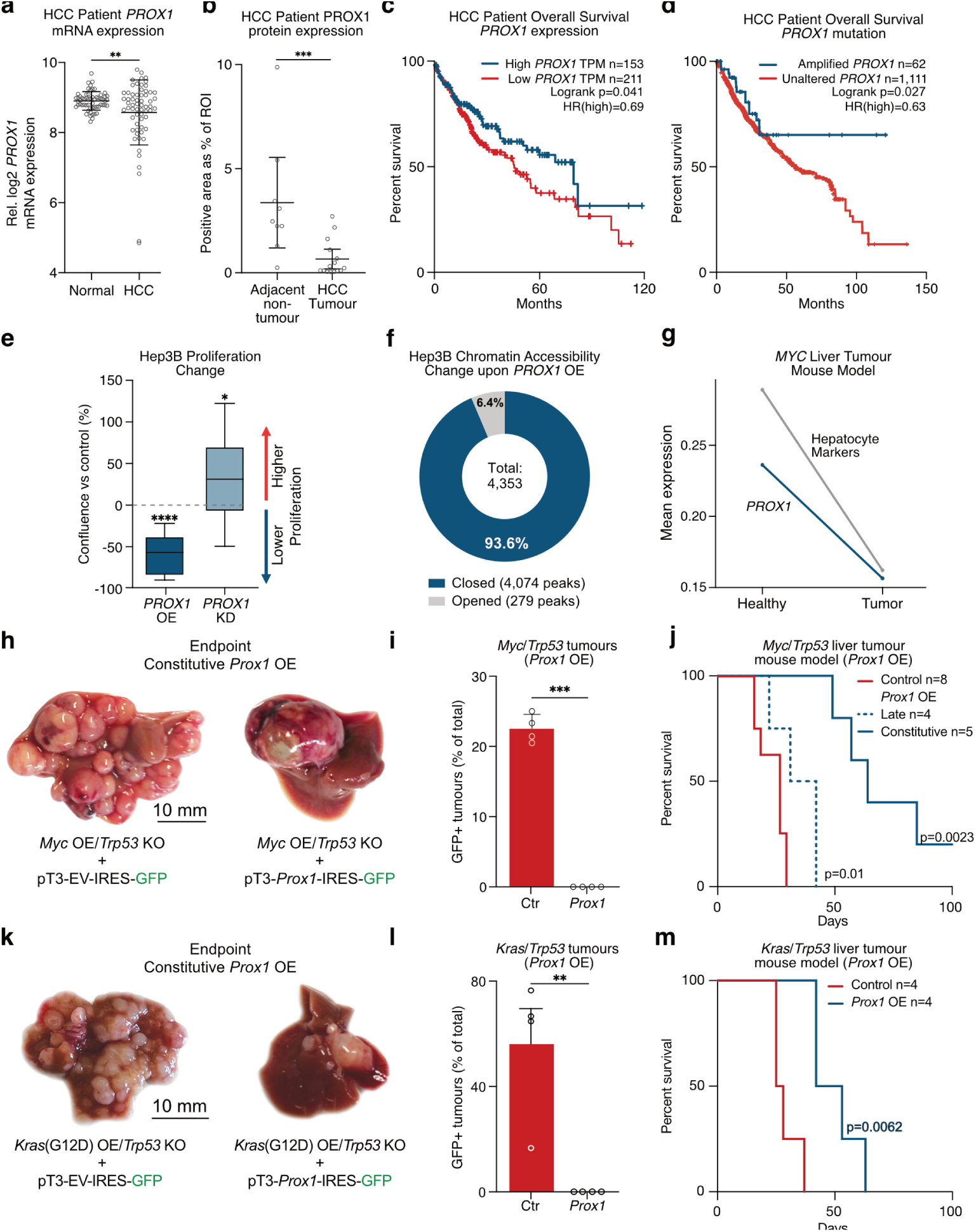
PROX1 suppresses hepatocyte transformation and liver cancer formation and progression. a, *PROX1* gene expression in tumour samples from HCC patients (n=62) and paired normal tissue (n=59) from the TIGER-LC dataset^27^. b, PROX1 protein levels in HCC patient liver sections within tumours and adjacent non-tumour tissues. c, Overall survival of 364 HCC patients segregated by *PROX1* expression levels (40% cutoff for high-expression cohort)^28^. d, Overall survival of 1,173 HCC patients segregated by *PROX1* mutation status (chromosomal amplification including *PROX1* or unaltered)^29–35^. e, Confluence percentage of Hep3B cells after 7 days of culture upon induction of *PROX1* shRNA-knockdown (KD) or overexpression (OE) normalised to uninduced controls. f, Differentially closed and opened regions two days upon *PROX1* overexpression in Hep3B cells compared to controls determined by ATAC-seq. g, *Prox1* and hepatocyte marker gene expression in healthy (day 0) and MYC-induced mouse HCC model (day 28) based on single-cell gene expression analysis^39^. Mean expression of Prox1 and hepatocyte signature genes across 7793cells is shown. h, Representative mouse livers following hydrodynamic tail vein injection (HDTVI) to induce *Myc* overexpression and *Trp53* knockout together with Constitutive *Prox1* overexpression (OE) (n=5). i, Percentage of GFP+ tumours in mice treated as in (h), indicating no *Prox1*-IRES-GFP positive tumours, at endpoint. j, Overall survival of mice treated as in (g) following Constitutive *Prox1* overexpression (OE) (n=5) or doxycycline-inducible Late *Prox1* OE (n=4) at day 14 compared to control (n=5 for Constitutive and n=3 for Late). k, HDTVI-induced livers tumours following *KrasG12D* overexpression and *Trp53* knockout together with Constitutive *Prox1* overexpression (OE) (n=4). i, Percentage of GFP+ tumours in mice treated as in (k), indicating lack of *Prox1*-IRES-GFP positive tumours. m, Overall survival of mice treated as in (k) following Constitutive *Prox1* or control overexpression (n=4). Bar graphs and scatter plots show mean values from specified biological replicates, error bars = SD, unpaired t-test (a, i and l), Mann-Whitney test (b), Log rank test (c,d,j, and m), and one-sample t-test assuming 0 as a theoretical mean (e), * p-adj < 0.05, ** p-adj < 0.01, *** p-adj < 0.001, **** p-adj < 0.0001.

### Dose-dependent suppression of cancer cell proliferation by PROX1

To investigate the effect of PROX1 levels on the physiology of human liver cancer cells, we manipulated its expression in an HCC cell line (Hep3B) *in vitro*. Specifically, we generated stable Hep3B cell lines with inducible *PROX1* overexpression or knockdown constructs, respectively (**Extended Data Figs. 2b,c; Methods)**. While *PROX1* overexpression decreased proliferation by 60%, shRNA-mediated depletion significantly enhanced proliferation *in vitro* (**Fig. 2e**). To investigate the molecular mechanism behind the antiproliferative activity of PROX1, we characterised the effects of PROX1 on chromatin organisation using an assay for transposase-accessible chromatin followed by sequencing (ATAC-seq). On day two of *Prox1* overexpression, we found 4,353 differentially-accessible peaks (padj < 0.05) of which 4,074 (93.6%) were closed compared to GFP control (**Fig. 2f**; **Extended Data Figs. 2d,e; Supplementary Table 3)**. To identify PROX1 target genes, we conducted CUT&RUN at the same time point with antibodies against PROX1 or IgG as control. We identified 16,183 peaks harbouring PROX1 motifs and found that PROX1 binding was enriched at sites that decreased in accessibility upon overexpression (Fisher test, p < 2.2e-16, odds ratio 1.6). The PROX1-bound regions included the MYC locus (**Extended Data Fig. 2f; Supplementary Table 4**). RNA-sequencing (RNA-seq) followed by Gene Set Enrichment Analysis (GSEA) and transcription factor importance analysis (GRaNPA; **Methods**) confirmed that *MYC* and MYC targets were downregulated upon *PROX1* overexpression, which coincided with increased expression of an apoptosis gene signature (**Extended Data Figs. 2f,g; Supplementary Table 5**). Next, we sought to investigate the dose-dependent effect of PROX1 levels on primary mouse liver cancer cells. To that end, we used lentivirus to introduce doxycycline-inducible *Prox1* in two primary cell lines derived from *in vivo* with distinct drivers, combining *Trp53* knockout with either *Myc* or *Kras*(G12D) overexpression^36^. In both models, we observed a significant reduction in proliferation in a PROX1 dose-dependent manner upon doxycycline titration (**Extended Data Figs. 2h-j**). This shows that PROX1 primarily closes chromatin in liver cancer cells and reduces their proliferation *in vitro* in a dose-dependent manner by regulating gene expression.

### PROX1 blocks hepatocyte transformation and liver cancer progression in mice

Next, we tested whether PROX1 can suppress liver tumourigenesis by preventing cell fate plasticity and tumour initiation in mice. First, we assessed the expression of *Prox1* using available single-cell data from an HCC mouse model^37^ and found that *Prox1* expression is lower in transformed cells compared to healthy hepatocytes (**Fig. 2g**). In addition, these tumour cells also exhibit decreased overall hepatocyte identity based on cell type-specific gene expression patterns (**Fig. 2g**). To investigate whether manipulating PROX1 can prevent liver tumour formation, we used an established HCC mouse model. To that end, we introduced stable *Myc* overexpression and *Trp53* knockout (*Myc*/*Trp53*) via transposable elements into mouse livers via hydrodynamic tail-vein injection (HDTVI) (**Fig. 2h**; **Extended Data Fig. 3a; Supplementary Table 6; Methods)**. After two weeks, mice developed multifocal liver carcinomas resembling HCC nodules with solid and trabecular growth patterns (**Extended Data Fig. 3b**). Tumours exhibited strong expression of the hepatocyte-specific transcription factor, hepatocyte nuclear factor-4α (HNF4α), did not express the biliary epithelial cell marker, keratin 19 (KRT19), and lacked glandular structures typical for cholangiocarcinomas (CCAs) (**Extended Data Fig. 3b**). Constitutive overexpression of *Prox1* in this model led to fewer tumour nodules at the endpoint (6.7 vs 39.5 nodules, p < 0.001) and, most strikingly, all tumours in the *Prox1*-IRES-GFP animals were GFP-negative, indicating that tumour cells strongly selected against *Prox1* overexpression (**Fig. 2i**; **Extended Data Figs. 3c-d**). Indeed, *Prox1* overexpression significantly increased median survival from 29 days to 64 days and caused a significant increase in overall survival duration with several mice surviving the 100-day experiment (**Fig. 2j**; **Extended Data Figs. 3l**). To test dose-dependent effects during *in vivo* tumourigenesis, we compared low with high PROX1 overexpression using PGK- and Ef1a-promoters, respectively, and found that only high PROX1 levels significantly increased survival (**Extended Data Figs. 3j-l**). This increase is biologically significant compared to other treatments such as the kinase inhibitor drug sorafenib, approved for the treatment of advanced primary liver cancer, which only increases survival in comparable mouse models by around 8 days^38^.

Given the striking benefits of *Prox1* overexpression on tumour formation, and because of the beneficial correlation between survival in HCC patients with *PROX1* amplification, which presumably occurs later in tumour development, we decided to also test the effect of PROX1 during tumour progression. To that end, we used the same HCC mouse model and combined it with doxycycline-inducible *Prox1* overexpression. *Prox1* expression was induced after tumour nodules had formed for 14 days post-HDTVI (**Extended Data Fig. 3e**). Two days following doxycycline treatment, we observed a >4-fold increase in apoptosis in tumours as judged by CASP3 histology compared to control (**Extended Data Figs. 3f-g**). These results were substantiated by gene expression analysis based on RNA-seq two days following *Prox1* overexpression, which showed upregulation of apoptosis and downregulation of pro-proliferative MYC gene signatures, mirroring the findings in Hep3B cells *in vitro* described above (**Extended Data Fig. 2g, Supplementary Table 7**). Importantly, late *Prox1* overexpression resulted in fewer GFP+ tumour nodules at the endpoint (219.5 vs 13.75 nodules, p < 0.001) and significantly increased median survival from 17 days to 36.5 days (**Fig. 2j**; **Extended Data Fig. 3h-i**), suggesting that PROX1 also blocks tumour progression. To determine whether PROX1 broadly suppresses liver cancer, we employed a second murine model induced by HDTVI-mediated oncogenic *Kras*(G12D) overexpression and *Trp53* knockout (*Kras*/*Trp53*) (**Extended Data Fig. 3m**). This model presents features of both HCC and CCA, characterized by morphologically complex tumours with loss of HNF4α and gain of KRT19 staining (**Extended Data Fig. 3n**). *Prox1* overexpression in this context also resulted in a significant reduction in tumour nodule count at the endpoint (3.25 vs. 19.5 nodules, p < 0.05), with all nodules lacking Prox1-IRES-GFP, and extended median survival from 26.5 days to 47.5 days (**Fig. 2k-m**; **Extended Data Figs. 3o-p**). Collectively, these findings demonstrate that PROX1 functions as a tumour suppressor by impeding tumour initiation and progression in distinct liver cancer models.

### PROX1 is required for efficient liver regeneration upon injury

Beyond development and cancer, cell fate plasticity also plays a key role in other fate transitions, such as during regeneration following injury or direct cell reprogramming. We therefore investigated whether PROX1 can also regulate cell fate plasticity in these processes. Upon liver injury, mature hepatocytes can undergo a dedifferentiation process associated with the reactivation of progenitor-like programs followed by proliferation and differentiation to regenerate functional hepatocytes^39^. Analysing available single-cell data along inferred pseudotime spanning pre-injury, injury and, post-injury cells from a 3,5-Diethoxycarbonyl-1,4-Dihydrocollidine (DDC)-induced mouse liver injury model^39^, we assessed the levels of *Prox1*, and found that its expression sharply decreased following injury and gradually recovered during hepatocyte regeneration **(Figs. 3a,b)**. Concordantly, hepatocyte identity decreased during the injury-induced dedifferentiation and was reactivated quickly during recovery. To test whether PROX1 is required during DDC-induced liver injury and regeneration we used conditional *Prox1* knockout mice^40^, in which exon 2 of *Prox1* is flanked by loxP sites (*Prox1^fl/fl^*) **(Fig. 3c)**. Compared to AAV Δcre-transduced controls, cre-transduced homozygous *Prox1^−/−^* deletion reduced the number and density of HNF4+ hepatocytes by ~33% and increased the number of KRT19+ cholangiocytes following regeneration **(Fig. 3d**; **Extended Data Figs. 4a-c)**. In addition, serum levels of the liver injury marker alkaline phosphatase (ALP) were elevated two-fold in *Prox1*-deleted mice after DDC withdrawal, while alanine aminotransferase (ALT) and aspartate aminotransferase (AST) showed the same trend but did not reach significance **(Fig. 3e**; **Extended Data Fig. 4d)**. Together, this suggests that PROX1 is required for efficient hepatocyte regeneration after liver injury.

**Fig. 3:**
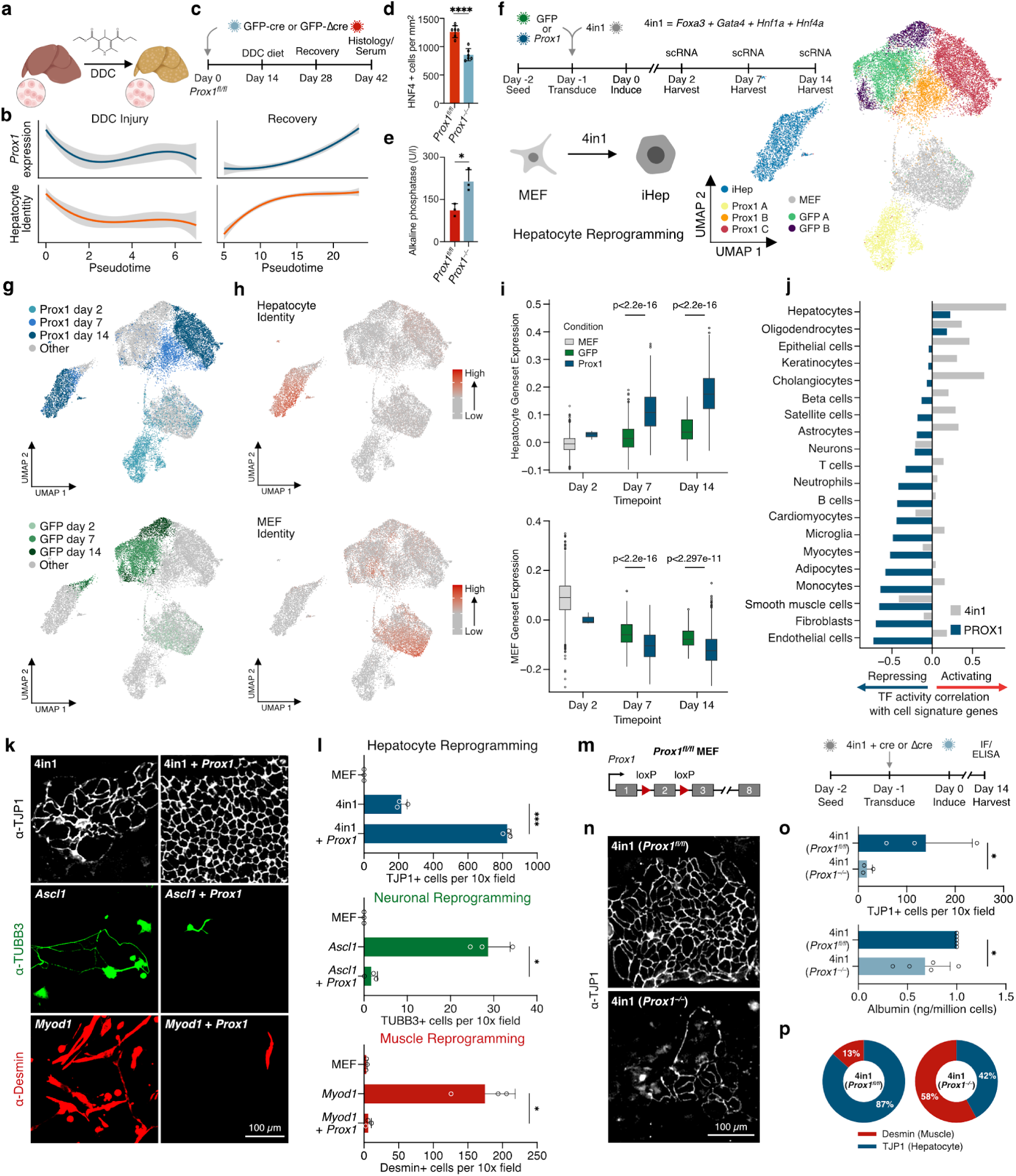
Multilineage repression by PROX1 is sufficient and necessary to promote hepatocyte cell fate. a, Schematic of DDC diet-induced liver injury and regeneration in mice. b, *Prox1* and hepatocyte signature levels of mice treated as in (a) based on single-cell gene expression pseudotime analysis across 1,866 cells ^39^. c, Liver injury and recovery as in (a) following cre-mediated deletion in conditional *Prox1^fl/fl^* knockout mice. d, Number of HNF4+ cells per area in the livers from mice treated as in (c) (n=3). e, Serum levels of Alkaline phosphatase from mice treated as in (c) (n=3). f, Hepatocyte reprogramming time course with or without *Prox1* analysed by single-cell RNA-seq, with 22,761 cells from two biological replicates following clustering and UMAP projection. g, Annotation of cells in (f) based on experimental treatment and time point. h, Projection of hepatocyte (top) and fibroblast (bottom) identity scores onto all cells in (f). i, Quantification of hepatocyte (top) and fibroblast (bottom) identity scores in hepatocyte cluster from (f), shown as boxplots with p-values (two-tailed t-test) for each time point and treatment. j, Correlation of various cell identity scores with 4in1 or PROX1 activity. k, Reprogramming of MEFs to induced hepatocytes (top), neurons (middle), or myocytes (bottom) with 4in1, *Ascl1*, or *Myod1* overexpression, respectively. Representative immunofluorescence of TJP1 (hepatocyte), TUBB3 (neuronal), or Desmin (myocyte) marker proteins at day 14 of respective reprogramming protocols with or without *Prox1* overexpression. l, Immunofluorescence quantification of cells in (k) (n=3). m, Knockout of *Prox1* during hepatocyte reprogramming via cre-mediated deletion of exon 2 in *Prox1^fl/fl^* MEFs. n, TJP1 immunofluorescence at day 14 of hepatocyte reprogramming in *Prox1^−/−^*or *Prox1^fl/fl^* cells. o, Analysis of cells in (n) quantifying the number of TJP1+ cells (n=3), and amount of Albumin secretion upon *Prox1* deletion (n=5). p, Proportion of reprogrammed cells in (n) positive for Desmin or TJP1. Bar graphs show mean values from specified biological replicates, error bars = SD, unpaired t-test, * p-adj < 0.05, *** p-adj < 0.001.

#### Liver cell fate is enhanced by active suppression of alternate cell identities

To experimentally investigate how PROX1 could enhance liver fate acquisition, we assessed the effects of *Prox1* overexpression during hepatocyte reprogramming. Specifically, we performed single-cell RNA-seq (scRNA-seq) of untreated MEFs and on days 2, 7, and 14 of 4in1-induced hepatocyte reprogramming together with overexpression of *Prox1* or GFP as control (**Fig. 3f)**. Following data processing and quality control filtering 22,761 cells from two reprogramming experiments were grouped into seven major clusters (**Fig. 3f**; **Extended Data Figs. 5a-d**). Cells with *Prox1* overexpression were clearly distinct from control cells and largely grouped into three clusters that closely corresponded to the three harvest time points (**Figs. 3f,g**). We identified two alternative fate clusters populated with GFP control cells (**Figs. 3f,g**). We scored each cell for expression of marker genes of different cell types from the curated Panglao database (**Supplementary Table 1; Methods**), and found one cluster strongly expressing MEF genes, reflecting the starting cell identity, and one cluster displaying a clear hepatocyte signature, indicating successful reprogramming (**Fig. 3h**). Overall, *Prox1* overexpression significantly increased the number of successfully-reprogrammed hepatocytes >7-fold (2,527 *Prox1* cells vs 347 control, p < 2.2e-16). Importantly, *Prox1* overexpression significantly increased hepatocyte cell identity in single cells at each time point, as measured by the expression of hepatocyte marker genes (**Fig. 3i**; **Extended Data Figs. 5e,f**). Strikingly, *Prox1*-overexpressing cells downregulated the initial MEF cell fate more efficiently and to a greater degree, and displayed lower expression of markers of alternate cell identities, such as fibroblasts, adipocytes, myocytes, and neurons (**Fig. 3i**; **Extended Data Figs. 5g-i**). These data indicated that PROX1 strongly enhances hepatocyte cell fate and suppresses gene expression programs of fibroblast and alternate cell types.

We next wanted to determine whether PROX1 promotes hepatocyte identity by regulating cell type-specific gene regulatory networks. To identify PROX1 target genes during reprogramming, we overexpressed FLAG-tagged *Prox1* or GFP and conducted CUT&RUN two days later with antibodies against FLAG or IgG as control (**Extended Data Fig. 6a**). We identified 25,519 peaks harbouring PROX1 motifs and found that PROX1 binding was enriched at gene promoters (**Extended Data Figs. 6b,c; Supplementary Table 4**). We defined genes within 1 kb from a PROX1 binding peak as targets and analysed their expression to determine PROX1 activity in single cells during reprogramming (**Supplementary Table 8; Methods**). PROX1 activity exhibited a significant negative correlation with the fibroblast identity score, suggesting that PROX1 directly represses genes of the donor cell fate (**Fig. 3j**; **Extended Data Fig. 6d**). Expanding this analysis to other cell fates, we found that PROX1 activity negatively correlated with all tested cell identities, except for hepatocyte and oligodendrocyte signatures (**Fig. 3j**). Interestingly, the activity of the liver inducers – FOXA3, GATA4, HNF1A, and HNF4A (4in1) – correlated positively with hepatocyte identity genes but also with several alternative cell identities, such as astrocytes, epithelial cells, and cholangiocytes (**Fig. 3j**). These signatures negatively correlated with PROX1 activity, suggesting that PROX1 can directly repress non-hepatic gene signatures, potentially activated by 4in1, to specifically promote the desired hepatocyte fate.

#### PROX1 can block alternate neuronal and muscle cell reprogramming

To determine whether PROX1 can actively repress alternative cell fates, we tested the effect of its overexpression on neuronal and myocyte reprogramming. Therefore, we overexpressed *Prox1* or GFP together with the pro-neuronal transcription factor *Ascl1*, which can convert fibroblasts into functional neurons^41^. Unlike reprogramming to hepatocytes, which was significantly enhanced by PROX1, neuronal reprogramming was almost entirely abolished upon *Prox1* overexpression, as determined by TUBB3 protein expression (**Figs. 3k,l**; **Extended Data Fig. 7**). Myocyte fate was among the top six signatures repressed by PROX1, and expression of the muscle marker protein Desmin was decreased by PROX1 during hepatocyte reprogramming **(Figs. 3j**; **Extended Data Fig. 7f)**. We therefore also tested whether PROX1 could actively suppress muscle cell reprogramming induced by *Myod1* overexpression^42^. PROX1 almost completely abolished induced myocyte reprogramming determined by Desmin protein expression and quantification of Desmin positive cells (**Figs. 3k,l**; **Extended Data Fig. 7**). Interestingly, while MYOD1 alone did not induce any liver-like cells, 18% of reprogrammed cells expressed TJP1 and displayed hepatic morphology upon *Prox1* co-expression (**Extended Data Fig. 7f**). Of note, the neuronal safeguard repressor MYT1L had an analogous effect, inhibiting muscle and liver cell reprogramming while promoting only neuronal identity during reprogramming (**Extended Data Figs. 7d-f**). This is in line with our previous finding that MYOD1 exhibits promiscuous activity and suggests that safeguard repressors, such as PROX1 or MYT1L, can redirect this activity to promote a specific, non-repressed cell fate^12^. In summary, PROX1 not only suppressed neuronal and muscle genes during hepatocyte reprogramming, but also strongly reduced, and even redirected, the cell fates induced by neuronal and muscle master regulators.

#### PROX1 is necessary to prevent alternate fates during hepatocyte reprogramming

Since exogenous *Prox1* overexpression was sufficient to suppress alternate fates, we sought to determine whether endogenous *Prox1* expression is necessary for efficient hepatocyte reprogramming. First, we performed shRNA-mediated *Prox1* knockdown during hepatocyte reprogramming. We confirmed *Prox1* downregulation by qRT-PCR, and found that several hepatocyte marker genes, such as *Alb*, *Krt18*, and *Cdh1*, were decreased upon *Prox1* depletion (**Extended Data Fig. 8**a). Monitoring Albumin secretion and TJP1 immunofluorescence confirmed significantly impaired liver reprogramming upon *Prox1* knockdown (**Extended Data Figs. 8b,c**). To substantiate these results, we prepared mouse embryonic fibroblasts from the conditional *Prox1* knockout mice (*Prox1^fl/fl^* MEF) (**Fig. 3m; Methods)**. As expected, cre-mediated homozygous *Prox1^−/−^* deletion significantly reduced the overall expression of *Prox1* and other liver markers compared to Δcre-transduced isogenic controls (**Extended Data Fig. 8d)**. Importantly, conditional *Prox1* deletion decreased the number of TJP1-positive hepatocyte-like cells >7-fold and significantly lowered Albumin secretion per cell (**Figs. 3n,o**). Conversely, the fraction of cells that expressed the muscle marker Desmin significantly increased upon *Prox1* deletion during liver reprogramming, while expression of E-cadherin decreased (**Fig. 3p**; **Extended Data Fig. 8e)**. Gene expression analysis by RNA-seq confirmed that genetic *Prox1^−/−^*deletion during hepatocyte reprogramming decreased the expression of many liver markers, and increased the expression of alternate fate markers such as neuronal *Map2* or muscle *Myh9* (**Extended Data Fig. 8f; Supplementary Table 10)**. Overall, these experiments showed that PROX1 is necessary and sufficient for efficient liver cell fate induction by repressing alternate fates.

#### Direct repression of PROX1 target genes enhances liver fate while activation induces fate plasticity

In mice, *Prox1* is also expressed in some neural stem cells in the hippocampus and cerebellum, in which it has been shown to promote neurogenesis^43,44^, and is also a key regulator of lymphatic endothelial cell differentiation and maintenance ^45,46^. In these contexts, PROX1 mainly activates gene expression in combination with coactivators, such as NR2F2 (a.k.a COUP-TFII)^47,48^. Conversely, in the liver, PROX1 was found to interact with corepressors, such as histone deacetylases^49^. To systematically expand on these findings we performed immunoprecipitation followed by mass spectrometry-based proteomics to identify PROX1 interaction partners in primary hepatocytes and neurons from mouse liver and hippocampus, respectively. Intriguingly, we found that PROX1 interacted with 9 out of 14 members of the repressive Nucleosome Remodeling and Deacetylase (NuRD) complex only in the liver. The NuRD complex contains histone deacetylases (HDACs) that catalyse the removal of acetyl groups from histones to mediate repression. This supports the notion that cell type-specific cofactor interactions enable PROX1 to switch between gene activation and repression (**Fig. 4a**; **Extended Data Fig. 9**). Hence, we wanted to uncouple cofactor-dependent effects and study which gene regulatory networks can be targeted directly by PROX1 to regulate cell identity. To directly activate or repress PROX1 target genes, we fused the DNA-binding domain (DBD) of PROX1 to either a transcriptional activator (VP64) or the Engrailed repressor (EnR) (**Fig. 4b)**. As a control, we expressed the DBD without an effector domain. While the DBD alone did not affect hepatocyte reprogramming, the repressor fusion improved hepatocyte fate induction similarly to full-length PROX1, as determined by the number of TJP1-positive hepatocyte-like cells and amount of Albumin secretion per cell (**Figs. 4c-e**; **Extended Data Figs. 10a-c**). Conversely, the activator fusion had a dominant negative effect and significantly impaired hepatocyte reprogramming. Furthermore, the fraction of reprogrammed cells that expressed alternate neuronal or myocyte markers was decreased by the repressor fusion and increased by the activator fusion (**Fig. 4f**; **Extended Data Fig. 10d**).

**Fig. 4:**
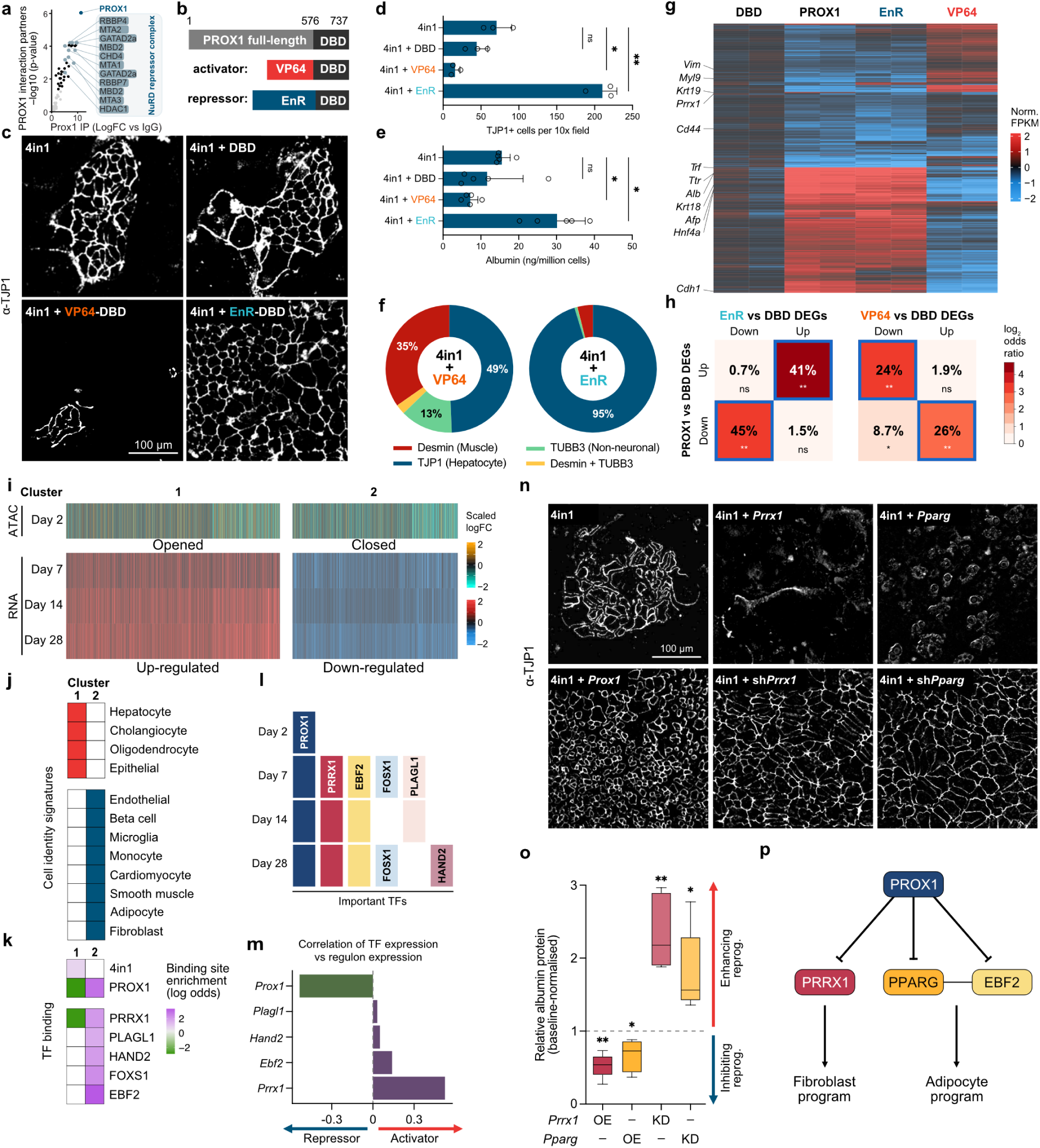
Alternative cell fate inducers are directly repressed by PROX1. a, PROX1 interaction partners identified by mass-spectrometry upon immunoprecipitation from mouse liver (n=4). b, Fusion proteins containing the PROX1 DNA-binding domain (DBD) and the VP64 activator or EnR repressor domains (not to scale). c, Representative TJP1 immunofluorescence upon 4in1-induced hepatocyte reprogramming with indicated PROX1 fusion constructs at day 14. d, Quantification of TJP1+ induced hepatocytes generated in (c) (n=3). e, Normalised Albumin secretion of cells in (c) (n=5). f, Proportion of reprogrammed cells in (c) positive for indicated cell type-specific markers and morphology. g, Normalised counts of differentially-expressed genes from RNA-seq of cells in (c), compared to DBD as a control, at day 7 (n=2). h, Percent overlap of up- and down-regulated genes in (g). Fisher test, * p < 1e-03, ** p < 1e-06. i, Chromatin accessibility (n=3) and gene expression analysis (n=2) comparing hepatocyte reprogramming with or without *Prox1* at indicated time points. Genes were clustered based on gene (red) up- and (bue) down-regulation and (yellow) increased vs (teal) decreased accessibility displayed as scaled logFC compared to control. j, Overlap of cell identity marker genes with genes present in each cluster. Overlaps with p-adj < 0.01 are shown. k, Enrichment or depletion of transcription factor binding from CUT&RUN (PROX1) or motif presence (all others) in the promoters of genes within each cluster. Log2 odds ratio with p-adj < 0.05 shown. l, Computational prediction of the transcription factors most important for PROX1-enhanced hepatocyte reprogramming over time. m, Expression correlation of indicated transcription factors and their target genes predicts activator vs repressor function. n, Representative TJP1 immunofluorescence upon 4in1-induced hepatocyte reprogramming with overexpression or shRNA-mediated knockdown of *Prrx1* or *Pparg* at day 14. o, Albumin protein quantification of cells in (n) by Western blot. Data is normalised to respective controls in boxplots (n=5). p, Proposed PROX1 gene regulatory network, by which repression of direct downstream transcription factors represses alternate cell identities. Bar graphs show mean values, error bars = SD, Dunnett’s test in (d and e), Fisher’s LSD test in (o), * p-adj < 0.05, ** p-adj < 0.01.

Since PROX1 inhibited neuronal and myocyte reprogramming, we also tested the effects of the activator and repressor fusions in these contexts. As expected, the repressor fusion reduced myocyte and neuronal induction, but a similar effect was observed when the DBD alone was expressed (**Extended Data Figs. 10a-c**). However, the fraction of TJP1-positive hepatocyte-like cells was increased upon repressor fusion expression during neuronal and myocyte reprogramming (**Extended Data Fig. 10d**). Conversely, the activator fusion reduced neuronal cell induction and enhanced myocyte induction, and increased the fraction of Desmin-positive cells in both settings relative to DBD (**Extended Data Figs. 10a-d**). Transcriptome analysis verified that, during hepatocyte reprogramming, the EnR fusion triggered a similar response in gene expression as full-length PROX1, causing induction of hepatocyte genes such as *Hnf4a* and *Krt18,* and concomitant repression of non-hepatocyte genes such as the cholangiocyte marker *Krt19* and the fibroblast transcription factor *Prrx1* (**Fig. 4g**; **Extended Data Fig. 10e; Supplementary Table 5**). Indeed, we found a significant overlap between genes downregulated by PROX1 and the repressor fusion (**Figs. 4g,h)**. Strikingly, the activator fusion had the opposite effect, decreasing hepatocyte gene expression and increasing alternate fate gene expression. Taken together, our experiments suggest that PROX1 directly binds and represses genes that drive non-hepatocyte fates, and thereby actively suppresses alternative fate trajectories to promote hepatocyte identity.

#### PROX1 decreases chromatin accessibility and expression of alternate fate signature genes

To investigate the effects of PROX1 on chromatin organisation during hepatocyte reprogramming, we performed ATAC-seq. On day two of 4in1-induced hepatocyte reprogramming, we found 111,411 differentially-accessible peaks (padj < 0.05) upon overexpression of *Prox1* compared to GFP control, of which 85,140 (76.4%) were closed (**Fig. 4i**; **Extended Data Figs. 11a,b; Supplementary Table 3**). We also overexpressed *Prox1* or GFP control without 4in1 and obtained a similar number of differentially-accessible peaks (109,917), of which 84,800 (77.1%) were closed upon *Prox1* overexpression. Hence, as in cancer cells (**Fig. 2f)**, PROX1 closed many chromatin regions, explaining ~80% of the variation between conditions (**Extended Data Figs. 11c,d**). Next, we characterised chromatin remodelling at PROX1-bound target genes using our PROX1 CUT&RUN DNA-binding data (**Extended Data Fig. 6; Supplementary Table 4**). We found a repressive signature and decreased accessibility at PROX1-bound sites upon *Prox1* overexpression, corroborating its repressive role (**Extended Data Fig. 11e; Methods**). Furthermore, 74% of genes with a PROX1-bound and differentially-accessible region in their promoters were downregulated upon *Prox1* overexpression, and none of the 71 upregulated genes were hepatocyte-specific markers (**Extended Data Fig. 11f**). This shows that PROX1 primarily closes chromatin at directly-bound target genes, reducing their expression in fibroblasts and the early stages of hepatocyte reprogramming.

To explore how PROX1 promotes hepatocyte fate at later stages of reprogramming, and to identify repressed PROX1 target genes that might mediate these effects, we performed a time course transcriptome analysis using bulk RNA-seq. Specifically, we assessed differential gene expression during hepatocyte reprogramming with GFP (control) or *Prox1* overexpression at days 7, 14, and 28 (**Fig. 4i)**. Globally, we found 8,036 protein-coding genes deregulated between *Prox1* and control at least at one of the three-time points (**Supplementary Table 5; Methods**).

Of these, 3,629 were up- and 3,264 were down-regulated consistently across all three-time points upon *Prox1* overexpression compared to control. Based on promoter accessibility at day two and differential gene expression over time, these genes were grouped into 2 clusters (**Fig. 4i)**. Genes in cluster 1 were upregulated throughout the 4 weeks and displayed increased promoter accessibility at day two upon *Prox1* overexpression (t-test, p < 1.487e-05) (**Extended Data Fig. 12a**). Cluster 1 was enriched for hepatocyte identity genes, but also contained some non-hepatic genes such as cholangiocyte and oligodendrocyte markers (**Fig. 4j**). On the other hand, cluster 2 was down-regulated upon *Prox1* overexpression throughout the 28-day reprogramming experiment and had reduced promoter accessibility at day two (t-test, p < 6.637e-09) (**Fig. 4i**; **Extended Data Fig. 12a)**. Genes in these clusters were enriched in many non-hepatocyte identity markers, including neuron and muscle genes, as well as fibroblast and adipocyte genes (**Fig. 4j**). During the 28-day reprogramming period, the expression levels of genes in cluster 2 increased in control cells but decreased upon *Prox1* overexpression (**Extended Data Fig. 12a**). Notably, genes from cluster 2 were expressed mostly in the PROX1-repressed alternative fate clusters of our single-cell dataset (**Extended Data Fig. 12b**). By contrast, genes from clusters 1 were highly expressed in reprogrammed hepatocytes in our single-cell transcriptomics dataset.

Next, we asked whether any of these clusters were directly regulated by our overexpressed transcription factors. To determine the direct effects of the liver reprogramming factors FOXA3, GATA4, HNF1A, and HNF4A (4in1), we retrieved the target genes of these four transcription factors and combined them into a 4in1 regulon containing 145 target genes (**Methods; Supplementary Table 8**). In addition, we defined a PROX1 regulon comprising 1,411 genes, which were bound by PROX1 based on CUT&RUN and showed direct regulation as determined through activator or repressor fusion transcriptome analysis (**Fig. 4g**; **Extended Data Fig. 6; Supplementary Table 8**). The hepatocyte gene-enriched cluster 1 was significantly enriched for 4in1 target genes and was strongly induced during reprogramming (**Figs. 4i-k**). Conversely, PROX1 target genes were significantly depleted in cluster 1, suggesting that 4in1 directly enhances hepatocyte maturation while PROX1 indirectly promoted this effect (**Figs. 4j,k**). Indeed, cluster 2, which contained 8 alternate fate signatures, was significantly enriched for direct PROX1 targets (**Figs. 4j,k**). Since this cluster was downregulated upon *Prox1* overexpression (**Fig. 4i**), this further supports that PROX1 mediates active repression of unwanted fates to promote liver cell induction and maturation.

#### *Prrx1* and *Pparg* are two alternate fate inducers repressed by PROX1

To understand how PROX1 silences many non-hepatocyte cell fates, and to identify key PROX1 target genes, we employed GRaNPA, a computational method to predict the importance of specific transcription factors based on differential gene expression^50^. We constructed gene regulatory networks for PROX1 and direct PROX1 targets annotated to be transcription factors. As expected, differential expression at day two is almost exclusively explained by PROX1 (**Fig. 4l**). At day 7, we predicted additional transcription factors, namely the direct PROX1 targets PRRX1 and EBF2, to regulate differential gene expression (**Fig. 4l**). Importantly, based on network analysis, PROX1 remained important at days 14 and 28, as did its direct targets PRRX1 and EBF2. The observation that PROX1 remains important throughout the 4-week experiment supports its function in hepatocyte fate maintenance (**Fig. 4l**). Starting from day 28, we found additional transcription factors predicted to be regulated by PROX1, such as the cardiac regulator HAND2^51^ (**Fig. 4l**). Strikingly, the regulons of all transcription factors targeted by PROX1 were enriched in the repressed non-hepatocyte cluster 2 (**Fig. 4j,k**). Unlike PROX1, all downstream transcription factors are predicted to be activators based on expression correlation with their target genes (**Fig. 4m**). Therefore, our gene regulatory network analysis suggests that PROX1 safeguards hepatocyte identity by silencing alternate cell fate genes both directly and by repression of transcription factors that activate non-hepatocyte gene programs.

The donor fibroblast identity and alternate adipocyte identity were among the gene signatures most strongly repressed by PROX1, based upon bulk and single-cell transcriptomic inference during hepatocyte reprogramming (**Figs. 3j and 4j**). In line with this, our analyses identified master regulators of both cell types as direct PROX1 targets. EBF2 is a co-activator of PPARG, and together they can drive adipogenesis^52,53^. PRRX1 is a master transcription factor of stromal fibroblasts, regulating the differentiation of mesodermal cell types and driving fibrosis during wound healing^54,55^. We verified that *Prrx1* and *Pparg* were bound by PROX1 at their promoters and displayed reduced chromatin accessibility upon *Prox1* overexpression, confirming that they are direct PROX1 targets (**Extended Data Fig. 13a**). We further overexpressed *Prrx1* or *Pparg* during 4in1-mediated hepatocyte reprogramming and observed, similar to *Prox1* deletion, reduced Albumin expression per cell by ~50% and impaired liver fate induction based on TJP1-immunofluorescence (**Figs. 4n,o**). In line, overexpression of *Prrx1* or *Pparg* together with *Prox1* cancelled the positive effects of PROX1 during liver reprogramming (**Extended Data Figs. 13b,c**). Conversely, shRNA-mediated depletion of *Prrx1* or *Pparg* increased Albumin expression per cell ~1-2 fold, and the number of TJP1-positive hepatocyte-like cells upon 4in1 expression (**Figs. 4n,o**; **Extended Data Figs. 13b,d,e**), similar to *Prox1* overexpression. Depletion of *Prrx1* or *Pparg* alongside *Prox1* overexpression did not further enhance hepatocyte reprogramming based on Albumin protein levels and TJP1-immunofluorescence, further indicating that both act downstream of PROX1 (**Fig. 4p**; **Extended Data Fig. 13e**). In summary, we find that PROX1 suppresses increased plasticity by repressing non-hepatic cell identities via direct transcriptional silencing of master regulators of alternate lineages, including fibroblast-specific *Prrx1* and the adipocyte regulator *Pparg*.

#### PROX1 regulates plasticity between cholangiocarcinoma vs hepatocellular carcinoma fates

Cellular plasticity and transdifferentiation also play important roles in liver cancer. The two dominant forms of primary liver cancer, cholangiocarcinoma (CCA) and hepatocellular carcinoma (HCC)^21,56^, which differ markedly in cellular composition and morphology^57,58^, can both arise from hepatocytes. However, the transcriptional mechanisms that regulate the transformation of hepatocytes to HCC vs CCA are largely unknown. To investigate whether PROX1 could play a role in this process, we assessed whether *PROX1* expression differs between CCA and HCC samples from 153 liver cancer patients^27^. We found that the median *PROX1* expression was 1.4-fold higher in HCC (n=62) patient samples than in CCA (n=91) (**Fig. 5a**). Interestingly, survival in HCC patients with high *PROX1* expression is significantly increased compared to CCA patients (**Extended Data Figs. 14a**). This was mirrored by the positive correlation of PROX1 expression and the HCC marker HNF4A (**Fig. 5b**). Intriguingly, not only the CCA marker KRT19 but also the PROX1 targets PRRX1 and PPARG exhibited a negative correlation with PROX1 expression in tumour tissue. This suggests that PROX1 could also prevent plasticity in primary liver cancer by suppression of alternate fate regulators and may regulate the transformation trajectory of hepatocytes towards HCC instead of CCA.

**Fig. 5:**
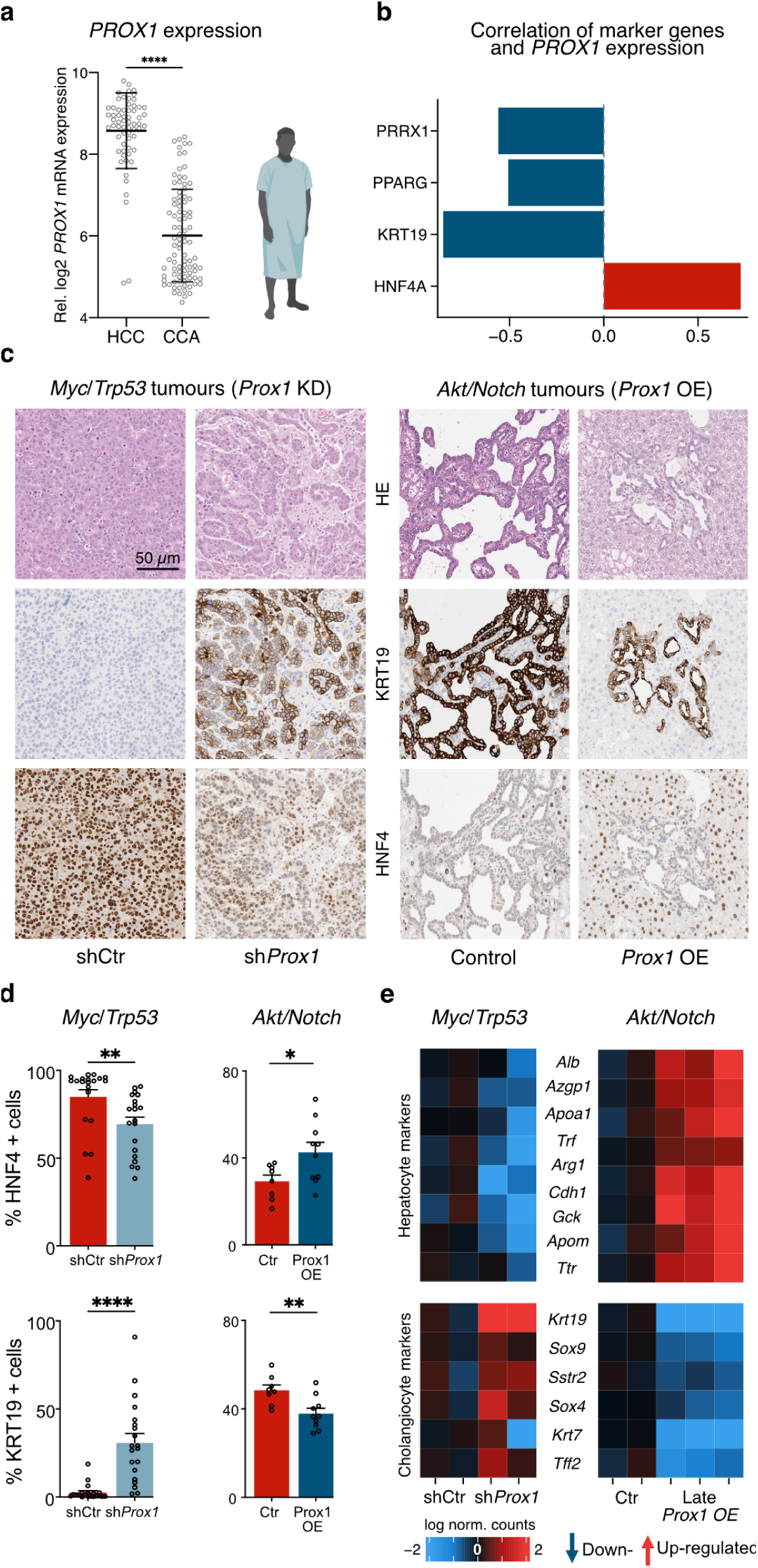
PROX1 regulates HCC to CCA fate trajectories in liver cancer. a, *PROX1* gene expression in tumour samples from cancer patients with HCC (n=62) and CCA (n=91) subtypes from the TIGER-LC dataset^27^. b, Correlation of indicated HCC and CCA markers and fate regulators with *PROX1* expression in patients from (a). c, Immunohistology of HDTVI-tumour models from HCC (*Myc*/*Trp53*) and CCA (*Akt*/*Notch*) mice at the endpoint generated using hematoxylin and eosin (HE) staining as well as KRT19 and HNF4 antibodies following *Prox1*-knockdown or overexpression compared to control (each n=5). d, Quantification of KRT19 (CCA-marker) and HNF4 (HCC-marker) positive cells in GFP+ tumours from (c) shown as percentage. e, Transcriptome analysis of tumour nodules from mice in (c) using RNA-seq following knockdown of *Prox1* (n=2) or Late *Prox1* overexpression (n=2-3). Heatmap of selected differentially expressed cholangiocyte- and hepatocyte-related genes is shown. Bar graphs and scatter plots show mean values from specified biological replicates, error bars = SD, unpaired t-test, ** p-adj < 0.01, *** p-adj < 0.001, **** p-adj < 0.0001.

To determine whether PROX1 loss influences liver cancer identity *in vivo*, we performed CRISPR-mediated *Prox1* knockout during HDTVI-mediated HCC formation in our *Myc*/*Trp53* model. Consistent with a tumour suppressor role, *Prox1* knockout in this model increased the number of tumours more than two-fold and decreased the median survival by 10 days (**Extended Data Figs. 14b-d**). To substantiate our results and directly follow perturbed cells, we performed knockdown of *Prox1* using shRNA constructs coupled to a GFP-reporter in the same model. *Prox1* knockdown led to a 10-fold increase in the number of microscopically detectable tumour nodules two weeks following HDTVI, but did not affect tumour number and survival at the endpoint (**Extended Data Figs. 14e-i**). Strikingly, *Prox1* knockdown induced a morphological shift from HCC towards CCA, with the formation of glandular tumour structures accompanied by significantly decreased HNF4 expression and concomitant increase in KRT19 expression (**Figs. 5c,d**).

Motivated by these findings, we investigated whether PROX1 gain could drive liver cancer identity from CCA towards HCC. To that end, we employed an well-established CCA mouse model using HDTVI-mediated overexpression of *Akt* and the *Notch1 receptor intracellular domain* (NICD) (*Akt*/*Notch*)^56^. After two to three weeks this model induced multifocal liver carcinomas with glandular structures and high KRT19 levels, which lacked HNF4α expression and mimicked CCA (**Fig. 5c**; **Extended Data Figs. 15a,b**). Constitutive overexpression of *Prox1* in this model led to fewer tumour nodules at the endpoint (59.8 vs 245.2 nodules, p < 0.05) and significantly increased median survival from 54 days to 88 days, indicating that *Prox1* overexpression also impaired hepatocyte formation also in this model (**Extended Data Figs. 15c-f**). Importantly, *Prox1* overexpression, when combined with *Akt*/*Notch,* induced a morphological shift from CCA towards HCC, with a reduction of glandular structures, significantly decreased KRT19 expression, and a concomitant increase in HNF4 expression (**Figs. 5c,d**). Strikingly, performing doxycycline-inducible late *Prox1* overexpression after 20 days of HDTVI-mediated tumour formation also decreased gland-like morphologies and reduced expression of the cholangiocyte markers KRT19 and SOX9 (**Extended Data Figs. 15g-j**).

Transcriptome analysis of tumour nodules from the CCA (*Akt*/*Notch*) and the HCC (*Myc*/*Trp53*) mice corroborated these shifts, with *Prox1* knockdown in HCC causing downregulation of several hepatocyte markers such as *Alb, Apoa1, Cdh1, Trf*, and *Ttr*, and upregulation of cholangiocyte markers, such as *Krt19* and *Sox9* (**Fig. 5e; Supplementary Table 7**), while *Prox1* overexpression in CCA resulted in the opposite effect. In line with expression in human liver cancer patient samples, we observed that manipulating PROX1 in mouse liver cancer models could switch the transformation trajectory of hepatocytes between CCA and HCC. Overall, this suggests that PROX1 can act as a safeguard repressor to maintain hepatocyte cell identity and prevent liver disease *in vivo*, with lower levels permitting increased plasticity and higher levels reducing the potential for transformation and transdifferentiation.

## Discussion

Several dozen transcription factors might be expressed in a given cell, but only a handful of so-called selectors or master regulators are sufficient to induce specific lineage identities by activating gene expression during development. Interestingly, ablating lineage regulators can cause expression of alternative lineage genes. For instance, the deletion of *Pax5* or *EBF1* in B cells results in the upregulation of myeloid gene expression ^59,60^. Similarly, genetic removal of *Bcl11b* in T cells leads to the derepression of NK cell-specific genes, facilitating their transdifferentiation into NK-T cells ^61^. Loss of some of these genes, such as *Pax5,* contributes to cancer formation^2,7,8^. In addition, mounting evidence suggests that repressive factors play critical roles in safeguarding cell fate induction and maintenance by preventing unwanted gene expression^10^. For instance, loss of repressive polycomb group proteins in mouse embryonic stem cells (mESCs) causes activation of developmental transcription factors^62^, and deletion of the BAF chromatin-remodelling complex member *Brm* during directed cardiogenesis from mESCs induces neuronal transcription factor expression, which shifts the identity of precardiac mesoderm to neural precursors^63^. Besides these ubiquitously-expressed repressive chromatin regulators, recent examples of cell type-specific transcription repressors have been reported to safeguard cell fate^10^. For example, the male germ cell-specific repressor Kmg inhibits the expression of somatic lineage genes in *D. melanogaster*^64^. Similarly, the neuron-specific repressor MYT1L prevents the expression of non-neuronal genes in neurons to induce and maintain mouse neuronal cell fate and function^12,13^. Here, we provide evidence that PROX1 represses non-hepatocyte genes to promote and cement liver cell identity.

Our findings allow us to outline a speculative model to explain the mechanism of action of such cell type-specific safeguard repressors. We propose that safeguard repressors regulate target genes that are inappropriately accessible^65^, and that they do so (i) based on their affinity to specific DNA motifs, while exhibiting (ii) cell type-specific, and (iii) continuous expression. Analysis of 1,296 transcription factors identified 27 potential safeguard repressor candidates across 18 cell types, with over 50%, including MYT1L, previously linked to promoting their predicted cell fates. PROX1, essential for hepatocyte commitment in mice and human cells^66–68^, meets the criteria of a hepatocyte safeguard repressor. First, PROX1 exhibits high lifelong expression in hepatocytes, enhanced reprogramming efficiency by 10-fold, and suppressed most tested alternate cell identity gene signatures. Second, deletion of Prox1 reduced hepatocyte reprogramming by 87% and impaired liver regeneration following injury. Third, PROX1 binding was associated with decreased chromatin accessibility and gene repression of alternate fate regulators.

This contrasts with reports studying the role of PROX1 in other lineages. Notably, in mice, *Prox1* is expressed in some cells in the hippocampus and cerebellum, in which it has been shown to promote neurogenesis^43,44^, and is also a master regulator of lymphatic endothelial cell differentiation^45,46^. In these contexts, PROX1 is reported to mainly induce gene expression in combination with coactivators, while in hepatocytes corepressor interactions are reported^47–49^. In line with this, we found that PROX1 was interacting with the repressive NuRD complex in the liver but not in the hippocampus. Decoupling PROX1 target gene regulation from the effects of recruited cofactors by fusion of the DNA binding domain of PROX1 to activator and repressor effector domains confirmed that PROX1 target gene repression promoted hepatic fate, while target gene activation induced alternate fates. Hence, depending on cofactor interaction a transcription factor can switch from a terminal selector into a safeguard repressor. Further studies are needed to characterise whether additional repressors guard cell fate canalisation in this manner, and if their life-long expression might help them to act as “terminal repressors” to prevent cell fate plasticity or even dedifferentiation and disease in mature cell types.

In line with this, we investigated the role of PROX1 in maintaining hepatocyte cell fate and preventing liver cancer *in vivo*. *PROX1* expression is decreased in liver cancer, and low levels are associated with poor survival in HCC patients, suggesting that PROX1 prevents hepatocyte plasticity and transformation. In mice, we find that *Prox1* overexpression strongly reduces neoplastic transformation and progression in a dose-dependent manner across three HDTVI-based liver cancer models. This contrasts with previous studies, in which *Prox1* overexpression in established tumour cells induced greater migratory and metastatic potential upon transplantation^25,69^. In line with a role in suppressing tumour initiation, *Prox1* knockdown or knockout accelerated tumour formation in our *in vivo* HCC model. Strikingly, resulting tumours displayed an identity switch away from HCC and towards CCA. Conversely, in CCA patients *PROX1* levels are low, and its overexpression in mouse models was able to shift CCA to HCC-like tumours. Indeed, hepatocytes are capable of giving rise to both HCC and CCA^21^, and our data indicate that PROX1 may be an important regulator of this decision, supporting the notion that safeguard repressors can prevent cell fate plasticity and block cancer development, progression and transdifferentiation *in vivo*.

In conclusion, our study shows that continuous cell type-specific repression of alternate fates is essential for cell fate induction and maintenance. Identifying and mechanistically characterising similar factors in other cell types, guided by computational tools such as the one presented here, could help generate cells for biomedical applications and reveal targets that prevent cell fate plasticity and disease.

## Methods

No statistical methods were used to predetermine the sample size for the experiments. Animals for primary cultures and *in vivo* experiments were selected randomly before indicated treatments. The investigators were blinded to the microscopy analysis and quantification. Otherwise, no blinding and randomisation were performed.

### Human material

Formalin-fixed, paraffin-embedded human liver tissue samples were retrieved from the Medical Faculty Mannheim, Heidelberg University, for immunohistological analyses. The study was approved by the local ethics committee (permit number 2012-293N-MA).

### Primary mouse cell lines

Mouse embryonic fibroblasts (MEFs) were harvested from E13.5 embryos of C57BL/6N (wild-type) or C57BL/6J (*Prox1^fl^*^/fl^) mice^40^ as described before^13,70^. The distal portions of all limbs from 3-4 embryos were dissected, placed in 100 μL trypsin, cut thoroughly, and incubated in a total of 1 mL trypsin (37°C, 15 min). Trypsin was inactivated by the addition of cell suspension to 25 mL MEF media (DMEM; Invitrogen) containing 10% cosmic calf serum (CCS; Hyclone), beta-mercaptoethanol (Sigma), non-essential amino acids, sodium pyruvate, L-glutamine, and penicillin/streptomycin (all from Invitrogen). MEFs were then cultured in MEF media and either cryopreserved or passaged twice using trypsin before reprogramming experiments.

### Hepatocellular carcinoma cell lines

Mouse primary liver cancer cell lines were derived from C57BL/6N female mice following HDTVI with either *Myc* OE/*Trp53* KO or *Kras*(G12D) OE/*Trp53* KO^36^, and human Hep3B cells were transduced with lentivirus prepared from indicated plasmids (**Supplementary Table 10**) in DMEM (Invitrogen) containing 10% foetal bovine serum (Sigma), non-essential amino acids, sodium pyruvate, L-glutamine, and penicillin/streptomycin (all from Invitrogen). Puromycin selection was performed (2 μg/ml) for 2 days to generate stable cell lines. Cellular proliferation rates were determined beginning 1 day after seeding with media containing 2 μg/ml or the indicated amount of doxycycline (Sigma) or without doxycycline (for controls) for a total of 7-10 days (depending on the proliferation rate of the line) using the IncuCyte S3 live-cell imaging system (Essen BioScience, Hertfordshire, UK). 4 brightfield images per well were acquired every 4 h at a magnification of 10× and analysed using the IncuCyte S3 2019B software. Experiments were performed in 2 to 3 biological replicates, with 5 to 6 technical replicates each.

### Animal experiments

For hydrodynamic tail vein injection (HDTVI), 2 mL of sterile 0.9% NaCl solution, corresponding to 10% of body weight, containing the plasmids of interest was injected into the tail vein of 8-week-old female C57BL/6N mice within 5 to 7 s^36,71–74^. Depending on the vector used, this technique allowed for liver-specific gene knockouts and/or overexpression by *in vivo* transfection of hepatocytes. Each mouse was injected with 20 µg of pX330-based plasmid for sgRNA-mediated gene knockout of *Tp53* or *Prox1,* and 10 µg of pT3-EF1a-based or pT3-PGK-based plasmid for transposon-mediated stable overexpression of *Myc, Kras, Akt or NICD*. For *Prox1* knockdown, 2 µg of CMV-Sleeping Beauty transposase and 20 µg of miR-E-based plasmid with *Prox1*-targeting shRNA and a GFP reporter were co-injected. For constitutive *Prox1* overexpression, 4 µg of CMV-Sleeping Beauty transposase and 10 µg of pT3-PGK-based or pT3-EF1a-based plasmid with *Prox1*-cDNA and a GFP reporter was co-injected (**Supplementary Tables 6 and 10**). For late *Prox1* overexpression, 4 µg of CMV-Sleeping Beauty transposase and 10 µg of pT3-Tre-based plasmid with *Prox1*-cDNA and a GFP reporter were co-injected. Each experimental group involving HDTVI contained at least 5 mice, with all mice monitored daily. For mice that received Tre-driven transgenes, a doxycycline-containing diet (6.25% doxycycline hyclate, Envigo Teklad, Indianapolis) was given beginning at day 14 or 20 post-HDTVI, depending on the model. Upon euthanasia of mice (at indicated time points or at humane endpoint), relevant organs were harvested and photographed. Survival data were analysed based on the time between HDTVI and euthanasia at a humane endpoint. After euthanasia, tumour samples were taken for RNA and protein analysis. The remaining tissue was incubated in 4% paraformaldehyde for a minimum of 24 h for subsequent histological analysis. For Diethyl-1,4-dihydro-2,4,6-trimethyl-3,5-pyridindicarboxylat (DDC, 137030 Sigma-Aldrich)-induced liver injury, 4-month-old *Prox1^fl^*^/fl^ mice^40^ were injected into the tail vein with 150 µl of PBS with 5 x 10^11^ genomic particles of Adeno-associated virus 8 (AAV8) carrying cre or Δcre-recombinase (**Supplementary Table 10)**. After 14 days, liver injury was induced by providing 0.1% DDC mixed with a standard diet (3437, KLIBA NAFAG) for 2 weeks followed by normal diet for another 2 weeks. After euthanasia, blood and tissue were harvested for subsequent analysis. All animal experiments were performed in compliance with ethical regulations and approved by the regional ethics board in Karlsruhe, Germany.

### Blood biochemical analysis

Blood was collected and serum was freshly isolated by centrifugation (12,000 g, 10 min, 4°C). Serum was stored at −20°C until analysis. Aspartate transaminase (AST), alkaline phosphatase (ALP), and alanine aminotransferase (ALT) levels were detected as per manufacturer instructions (Fujifilm DRI-CHEM SLIDE).

### Histology

Paraformaldehyde-incubated livers were embedded in paraffin and 2 μm slices were processed as previously described for immunohistochemistry (IHC) staining^75^. BOND-MAX (Leica Biosystems) was used for automated staining. BondTM citrate solution (AR9961, Leica), BondTM EDTA solution (AR9640, Leica), or BondTM proteolytic enzyme kit (AR9551, Leica) were used for antigen retrieval (Supplementary Table 15). Sections were incubated in antibodies diluted in BondTM primary antibody diluent (AR9352, Leica Biosystems) followed by secondary antibody (Leica Biosystems) incubation and staining with Bond Polymer Refine Detection Kit (DS9800, Leica Biosystems). Slides were scanned with an Aperio AT2 slide scanner (Leica Biosystems) at 20x, then annotated and analysed with Aperio ImageScope (v12.4.0.5043, Leica) for determining the size of tumour nodules. Marker staining quantifications were analyzed in QuPath (v0.4.3)^76^ using the positive cell detection option with the same settings for each marker across all sections. Certified pathologists (H.W. and D.T.) performed the histopathological analysis of paraffin-embedded liver tumour sections.

### PROX1 immunoprecipitation and LC-MS/MS analysis

For each immunoprecipitation, one hippocampus or liver of 2 to 3-month-old mice was used per biological replicate. The fresh tissue was lysed in 1 mL lysis buffer containing (in mM): 0.5% Tween-20, 50 Tris pH 7.5, 2 EDTA, 1 DTT, 1 PMSF, 5 NaF (all from Sigma), and complete protease inhibitor (Roche) for 15 min at 4 °C and processed for immunoprecipitation as described previously^13,15^ using 2 μg PROX1 or control IgG (Sigma) antibody per reaction (**Supplementary Table 11**). After elution bound proteins were enzymatically digested with trypsin using an AssayMAP Bravo liquid handling system (Agilent technologies) running the autoSP3 protocol as described here^77^. A LC-MS/MS analysis was carried out using a Vanquish Neo UPLC system (Thermo Fisher Scientific) directly connected to an Orbitrap Exploris 480 mass spectrometer for a total of 60 min per sample. Peptides were online desalted on a trapping cartridge (Acclaim PepMap300 C18, 5µm, 300Å wide pore; Thermo Fisher Scientific) with a loading volume of 60 ul using 30 ul/min flow of 0.05% TFA in water. The analytical multistep gradient (300 nl/min) was performed with a nanoEase MZ Peptide analytical column (300Å, 1.7 µm, 75 µm x 200 mm, Waters) using solvent A (0.1% formic acid in water) and solvent B (0.1% formic acid in acetonitrile). For 45 min the concentration of B was linearly ramped from 5% to 30%, followed by a quick ramp to 80%, after four min the concentration of B was lowered to 2% and a 3 column volumes equilibration appended. Eluting peptides were analyzed in the mass spectrometer using data-dependent acquisition (DDA) mode. A full scan at 60k resolution (380-1400 m/z, 500% AGC target, 100 ms maxIT) was followed by up to 1.5 s of MS/MS scans. Peptide features were isolated with a window of 1.2 m/z, fragmented using 26% NCE. Fragment spectra were recorded at 15k resolution (100% AGC target, 150 ms maxIT). Dynamic exclusion was set to 10 s. Each sample was followed by a wash injection to avoid carry over. System readiness was assessed before, during and after the measurements via an in-house QC pipeline. Data analysis was carried out by MaxQuant (version 2.1.4.0)^78^ using an organism-specific database extracted from Uniprot.org (mouse reference database with 1 protein sequence per gene, containing 21,957 unique entries from May 3rd, 2023). Settings were set to default with the following adaptions. Separate parameter groups were assigned for liver and hippocampus samples. Separate Label free quantification (LFQ) per parameter group was enabled. Besides the LFQ approach based on the MaxLFQ algorithm^79^, quantification was also done based on iBAQ-values^80^. The statistical analysis of proteins has been conducted as follows: Adapted from the Perseus recommendations^81^, protein groups with valid values in 70% of the samples of at least 1 condition were used for statistics. In addition, missing values, being completely absent in 1 condition, were imputed with random values drawn from a downshifted (2.2 standard deviation) and narrowed (0.3 standard deviation) intensity distribution of the individual samples. For missing values with no complete absence in one condition, the R package missForest^82^ was used for imputation. No additional normalization was applied to the iBAQ values that were used in the statistical analysis. The statistical analysis was performed with the R-package limma^83^ with an adapted contrast setup from chapter 9.5 Interaction Models. Within the eBayes function the options robust and trend were set to TRUE. The p-values were adjusted with the Benjamini–Hochberg method for the multiple testing. PROX1 interactors were considered significant with an absolute LogFC > 1, p-value < 0.05, and quality score > 0.5. In addition, interactors were filtered according to nuclear location (based on https://www.proteinatlas.org/about/download). The mass spectrometry data analysis can be found in **Supplementary Table 9**.

### Recombinant virus production

Lentivirus was produced through transfection of lentiviral backbones containing indicated transgenes along with third-generation packaging plasmids into HEK293T cells according to the Trono laboratory protocol (**Supplementary Table 10**)^84^. Lentivirus was concentrated from HEK293T culture supernatant through ultra-centrifugation (23,000 rpm, 2 h, 4°C) and stored at −80°C or used immediately. AAV8 vectors were produced by polyethyleneimine (PEI) triple-transfection of HEK293T cells using indicated plasmids as described before (**Supplementary Table 10**)^85^. AAV vectors were purified using iodixanol gradient density centrifugation followed by buffer exchange to PBS. AAV vector quantification was conducted by droplet digital (dd)PCR using the BioRad ddPCR system. Each 20 µL PCR contained 5 µL diluted virus template, 10 µL of the ddPCR Supermix for Probes (no dUTP; BioRad), 4 µL nuclease-free H2O and 1 µL ITR-primer/probe mix (final concentration: 900 nM for primers, 250 nM for probes; ITR_f: GGAACCCCTAGTGATGGAGTT, ITR_r: CGGCCTCAGTGAGCGA, ITR_probe: HEX-CACTCCCTCTCTGCGCGCTCG-BHQ1). The measured copy number of vector templates per reaction was corrected by the input volume and dilution factor to calculate vector genomes per µL vector stock.

### Direct reprogramming from MEFs

Wild-type or *Prox1^fl^*^/fl^ MEFs were transduced by incubation with lentivirus prepared from indicated plasmids (**Supplementary Table 10**) in MEF medium with 8 μg/ml polybrene (Sigma) for 16 to 20 h. Medium was exchanged to MEF medium containing 2 μg/ml doxycycline (Sigma) to induce transgene expression. For myocyte and neuronal reprogramming^70^, MEFs were transduced with lentivirus containing *Myod1* or *Ascl1,* respectively, along with rtTA-containing lentivirus. After 48 h, all medium was exchanged with N3 medium (DMEM/F12) containing N2 supplement, B27, 20 μg/ml insulin, penicillin/streptomycin (all from Invitrogen), and doxycycline to continue transgene expression. Medium was changed every 2 days for the remainder of the reprogramming. For hepatocyte reprogramming^23^, MEFs were seeded onto collagen-coated plates and transduced 1 day later with 4in1-containing lentivirus. 1 day following doxycycline induction the medium was supplemented with 0.5× volume Hepatocyte Culture Medium (HCM, Lonza) containing 5% FBS (Life Technologies) and 2 μg/ml doxycycline. On day 2, all medium was exchanged to a mixture of ⅓ MEF medium and ⅔ HCM + 5% FBS with 2 μg/ml doxycycline. On day 3, all medium was exchanged to HCM + 5% FBS with 2 μg/ml doxycycline. Medium was then changed every 2 days for the remainder of reprogramming. For domain fusion experiments we followed published protocols^70,86^, and for shRNA-based knockdown we treated cells with lentivirus targeting the indicated gene with 2 independent-shRNA constructs (**Supplementary Table 10**).

### Immunofluorescence quantification

To calculate the efficiency of neuronal induction, the total number of TUBB3-expressing cells with complex neurite outgrowth (cells with a round cell body and at least one thin process with a length at least double the diameter of the cell body) was counted manually^13^. Any TUBB3-positive and Desmin-negative cells that did not meet the morphological criterion were considered TUBB3-positive, non-neuronal cells (**Extended Date Fig. 6f**). To calculate the efficiency of myocyte cell induction, the total number of Desmin-expressing cells was counted manually. Dual TUBB3- and Desmin-positive cells were considered as a separate category of mixed identity cells. To calculate the efficiency of hepatocyte reprogramming, the total number of cells for which TJP1 staining formed a complete border around the nucleus (stained by DAPI) was counted manually. All quantifications were performed 14 days after transgene induction by immunofluorescence microscopy. Fluorescence micrographs were captured automatically using a Nikon Ti2 microscope with a Ti-HCS system, the Nikon S Plan Fluor ELWD 20× NA 0.45 objective, Nikon DS-Qi2 CMOS camera (2404×2404), and Lumencor Sola SE II light source. Quantifications were based on the mean number of positive cells across 10 randomly selected 20× magnification fields of view per biological replicate, with at least 3 biological replicates. The number of reprogrammed cells in each treatment condition was then normalised to the number of reprogrammed cells in the control condition. To calculate the fraction of total reprogrammed cells positive for indicated markers and morphological criteria, the number of cells in each of the 5 categories was divided by the total number of cells present in any of the 5 categories, pooled across all replicates.

### Computational safeguard repressor screen

Using single-cell gene expression and cell type annotations from Tabula Muris^87^, we analysed the expression of 1,296 transcription factors^88^ in 18 selected cell types using median normalised CPM units. In addition, we retrieved the top 1,000 cell type-enriched genes using Seurat FindMarkers() (**Supplementary Table 1**). In the promoters of each of these signature genes (±2 kb around the TSS), we determined the number of DNA-binding motifs for each transcription factor (derived from CIS-BP 1.94d^88,89^ mapped to mm10). Pairwise comparisons were performed between cell-type-specific gene signatures to remove overlapping genes and calculate mean transcription factor motif density. For each cell type, transcription factor expression specificity and transcription factor binding motif enrichment in signature gene promoters were calculated by Z-score scaling to obtain Z_expression_ and Z_motif_ respectively. The higher Z_expression_, the more specifically the transcription factor is expressed in the corresponding cell type. Transcription factors with positive Z_motif_, and high Z_expression_ might function as activators of cell type-specific genes. Conversely, a negative Z_motif_, indicates depletion of DNA binding motifs of this transcription factor at genes specific to the analysed cell type, indicating that it could act as a safeguard repressor by silencing cell type-unspecific genes. We harnessed our PROX1 CUT&RUN peaks, filtered using Hocomoco v12 motifs ^90^, to assess the gene signature binding enrichment using a one-tailed hypergeometric test. In addition, we calculated a safeguard repressor score for each transcription factor defined as the sum of Z_expression_ and –1 * Z_motif_, following Z_max_-normalisation to ensure equal weighting of expression and motif bias (**Fig. 1b**; **Extended Data Fig. 1c; Supplementary Table 2)**. Transcription factors were further annotated as cell type-specific and life-long expressed using data from Tabula Muris Senis^17^ based on the following two criteria: (i) the mean expression in Tabula Muris Senis must be at least 50% of that in Tabula Muris; (ii) the mean expression must be higher in a specific cell type compared to the mean expression across all 18 cell types in Tabula Muris. In addition, we categorised TFs as life-long expressed using a mouse developmental gene expression atlas ^18^ for the brain, heart, and liver if they exhibited high expression (in the highest quantile) in a continuous manner (in more than 70% of developmental time points and replicates). We analysed safeguard candidates for microglia, neurons, oligodendrocytes, and astrocytes from the brain data; cardiomyocytes from the heart data; and hepatocytes from the liver data. 77% of the safeguard repressor candidates in these cell types (17/22), including *Prox1* and *Myt1l*, were found to be life-long expressed in an organ-specific manner, which compared to all TFs expressed in these tissues reached statistically significant enrichment using a Fisher’s exact test. Top safeguard repressor candidates were also categorised as activators, repressors, or dual activator/repressors as well as having a known tumour suppressor role or not, in the cell type of interest, based on literature review (**Supplementary Table 2**).

### Bulk RNA-seq library generation

Primary mouse liver tumour samples were dissected and placed into TRIzol (Invitrogen). Samples were crushed and then homogenised using QIAshredder (Qiagen). Reprogrammed hepatocytes or Hep3B cells were harvested from culture plates by the addition of TRIzol to the cultures at indicated time points. RNA harvested in TRIzol was isolated using the RNA Miniprep kit (Zymo Research). For bulk RNA sequencing, libraries were prepared according to the dUTP protocol^91^ and paired-end sequencing (2×100bp) was performed on the NovaSeq 6000 or NovaSeq X Plus platforms (Illumina).

### RNA-seq data processing

Raw reads were mapped to the reference genome mm10 or hg38 using STAR^92^. Differential gene expression was determined using DESeq2 (R package version 1.28.1)^93^ with size factor normalisation and Wald significance tests. For bulk MEF reprogramming data, we used the primary MEF line as a covariate. ComplexHeatmap (version 2.12.1)^94^ was used to generate heatmaps. In **Fig. 4i**, genes were included in the analysis if they had an abs(log2FC) > 0.75 at any of the time points.

### CUT&RUN library preparation

Cells were harvested with Accutase and strained through a 70 μm strainer followed by CUT&RUN processing^95^. 250,000 cells were washed twice with 1 mL wash buffer (20 mM HEPES-KOH pH 7.5, 150 mM NaCl, 0.5 mM spermidine, and 1X Roche Complete Protease Inhibitor), then resuspended in 200 μL of ice-cold cell lysis buffer (10 mM Tris-HCl pH 7.5, 10 mM NaCl, 3 mM MgCl_2_, 0.1% Tween-20, 0.1% NP-40, and 1% BSA in ddH2O) for 3-5 min. 1 mL of ice-cold wash buffer was then added to stop cell lysis, and nuclei were centrifuged (500g, 10 min, 4°C). Nuclei were resuspended in 200 μL wash buffer, and 100,000–150,000 nuclei were taken to another tube. Concanavalin-A beads (Polysciences) were pre-activated in cold binding buffer (20 mM HEPES-KOH pH 7.5, 10 mM KCl, 1 mM CaCl2, and 1 mM MnCl2). Nuclei were centrifuged and buffer was removed. Activated beads were then added to the pellet. The bead-cell suspension was rotated (RT, 10 min). The supernatant was removed on a magnet and the beads were resuspended in antibody buffer (0.2 mM EDTA, 0.05% w/v digitonin in wash buffer). 1 µg primary antibody (rabbit anti-FLAG, *in vitro*, or rabbit anti-Prox1, *in vivo*) or control (mouse IgG, Sigma) was added (**Supplementary Table 11**), and cells were incubated on a nutator (overnight, 4°C). Beads were washed twice in digitonin-wash buffer (0.05% w/v digitonin in wash buffer), resuspended in 700 ng/mL pAG-MNase (Protein Expression and Purification Core Facility, EMBL, Heidelberg) in digitonin-wash buffer, and rotated (4°C, 1 h). pAG-MNase-loaded beads were then washed twice in digitonin-wash buffer, resuspended in digitonin-wash buffer, and placed on ice. 1 uL of 100 mM CaCl2 was added to induce chromatin digestion, and the mixture was incubated on ice (30 min). 50 uL of 2x stop buffer (340 mM NaCl, 20 mM EDTA, 4 mM EGTA, 0.05% w/v digitonin, 50 ug/mL RNase A, 50 ug/mL glycogen, 0.5 ng/mL spike-in E. coli DNA) was added, and the suspension incubated at 37°C for 10 min to release chromatin fragments from cells. The supernatant was subjected to phenol-chloroform extraction, and purified DNA fragments were used for library preparation with NEBNext DNA Library Prep Kit for Illumina (NEB E7645). Libraries were then sequenced (paired-end, 2×40 bp) on the NextSeq 550 and 2000 platform (Illumina). Livers from 2 to 3-month-old C57BL/6J mice were harvested and directly processed for nuclei isolation using liver swelling buffer (10 mM Tris pH 7.5, 2 mM MgCl2, 3 mM CaCl2) with the help of a douncer. Homogenized tissue was passed through a 70 µm strainer and centrifuged (400g, 5 min, 4°C). Tissue pellets were resuspended in liver lysis buffer (10 mM Tris pH 7.5, 1% NP-40, 2 mM MgCl2, 10% Glycerol, 3 mM CaCl2) and centrifuged again. Pellets containing the nuclei were washed twice in PBS and then the same protocol as for cells was followed. Hippocampi from 2 to 3-month-old C57BL/6J mice were harvested and directly processed for nuclei isolation. Tissue was lysed in brain lysis buffer (0.5% Tween-20, 50mM Tris pH 7.5, 2mM EDTA, 1x DTT, 1x NaF, 1x Protease inhibitor cocktail) by mechanical homogenization in a 1.5 mL Eppendorf tube. After 15 min of incubation on ice, nuclei were pelleted by centrifugation (1 min, 3200 rpm, 4°C). Then, the samples were processed as described for cells. For mouse livers and hippocampi, 500,000 nuclei were used per sample instead of 100,000-150,000 for cells. In addition, 5 μg of target antibody was used per sample.

### CUT&RUN data processing

CUT&RUN data were analysed using the nf-core/cutandrun pipeline v1.0.0 with Nextflow version 21.05.0^96,97^. Reads were aligned to mm10 or hg38. Software versions: bedtools (v2.30.0)^98^, bowtie2 (2.4.2)^99^, deeptools (v3.5.0)^100^, DESeq2 (v1.28.0)^93^, fastqc (v0.11.9), multiqc (v1.11)^101^, picard (v2.23.9)^102^, python (v3.8.3), samtools (v1.10)^103^, Genrich (v0.6.1)(https://github.com/jsh58/Genrich), TrimGalore (v0.6.6)^104^, ucsc (v377). Consensus peaks were defined by running Genrich -m 30 -e chrM -r -l 5 -q 0.3. To determine peaks containing motifs, we mapped the PROX1 binding motif (CIS-BP ID: M03445_2.00) onto mm10 or hg38, extended each resulting motif peak to a total width of 50 bp, and ran bedtools intersect -wa on the CUT&RUN consensus peaks and motif peaks, respectively. Genes defined as direct target genes based upon CUT&RUN were determined by running Homer annotatePeaks.pl on the final peak set and filtering for genes with a peak within ±1kb of their TSS. An overview of the CUT&RUN experiments is in **Supplementary Table 13**, and the final motif-containing peaksets for liver, Hep3B, and induced hepatocytes are in **Supplementary Tables 4**.

### ATAC-seq library preparation

Nuclei from liver and hippocampus tissues were isolated as described for CUT&RUN. Cells on culture plates were washed twice with PBS and detached by Accutase digestion (4-6 min, RT) followed by ATAC processing. Cell suspensions were placed into an equal volume of MEF medium followed by centrifugation (500g, 5 min, 4°C) before resuspension in ice-cold PBS + 1% BSA. Cells were then strained through a 70 μm filter, and centrifuged (500g, 5 min, 4°C). 50,000 cells or nuclei were resuspended in 50 μL of ice-cold cell lysis buffer containing 0.1% NP-40 and 0.01% digitonin in wash buffer (10 mM Tris-HCl pH 7.5, 10 mM NaCl, 3 mM MgCl_2_, 0.1% Tween-20, and 1% BSA in ddH2O) for 3-5 min. 1 mL of ice-cold wash buffer was then added to stop cell lysis, and nuclei were centrifuged (500g, 10 min, 4°C). 4.5 μL of each of the tagmentation oligos, Tn5_ME and Tn5-R1N (**Supplementary Table 12**), were annealed in oligo annealing buffer (10 mM Tris-HCl pH 7.5, 50 mM NaCl, and 10 mM EDTA final concentration in ddH2O) by heating at 95°C (3 min) followed by a ramp down by 1°C to 25°C. The same procedure was performed for Tn5_ME and Tn5-R2N (**Supplementary Table 12**). Tn5 was assembled with annealed oligos by combining 50 μL Tn5 (1 mg/mL stock) with 25 μL annealed Tn5_ME and Tn5-R1N and 25 μL Tn5_ME and Tn5-R2N. Nuclei were resuspended in 40 μL tagmentation buffer (38.8 mM Tris-acetate, 77.6 mM K-acetate, 11.8 Mg-acetate, 18.8% dimethylformamide, and 0.12% NP-40 in ddH2O), to which 5 μL ice-cold PBS + 1% BSA and 5 μL pre-assembled Tn5 was added. Samples were incubated on a Thermomixer (37°C, 30 min, 500 rpm) before being subjected to MinElute (Qiagen) cleanup to extract tagmented DNA. Eluted DNA was pre-amplified with P5 and P7 primers (**Supplementary Table 12**) using the NEBNext HF 2x PCR Master Mix (New England Biolabs) in a thermocycler set to 72°C (5 min), 98°C (30 sec), and 5 cycles of 98°C (10 sec), 63°C (30 sec), and 72°C (1 min). A qPCR side reaction was performed with the resulting pre-amplified libraries to determine the necessary additional cycles (5 cycles fewer than the number of cycles corresponding to ⅓ of max fluorescence) for complete amplification. After finishing amplification, 50 μL of each library was subjected to 2-sided size selection by addition of 27.5 μL (0.55x) AMPure XP beads, incubation (5 min), transfer of supernatant to new tubes, addition of 42.5 μL (1.4x) AMPure XP beads to the supernatant, incubation (5 min), three washes with 80% ethanol, and elution of DNA. Resulting libraries were then sequenced on the NextSeq 2000 platform (Illumina).

### ATAC-seq data processing

ATAC-seq data from *in vitro* samples were analysed with a custom Snakemake pipeline. Raw reads were quality-checked with fastqc (v0.11.8), trimmed with trimmomatic (v0.38), and aligned to UCSC mm10 or hg38 with bowtie2 (v2.3.4.3)^99^. Aligned reads were cleaned and base-recalibrated (to take account of Tn5 insertion biases) with samtools (v1.10)^103^ and picard (v2.18.16)^102^. Reads were filtered with bedtools (v2.27.1)^98^, samtools, and picard. Peaks were called using macs2 (2.1.2) and coverage was calculated with deeptools (v3.1.3)^100^. Final quality checks were performed with multiqc (v1.6)^101^. Differential peak analysis was performed with DiffBind (v3.4.11)^105^ as described in the authors’ vignette (same version). ATAC-seq from *in vivo* samples were analyzed using the nf-core/atacseq pipeline v2.1.2 with Nextflow version 23.10.1. Reads were aligned to mm10. Consensus peaks were defined by running Genrich -j -m 30 -e chrM -r -l 15 -q 0.01 -a 500 -g 15 -l 15 -d 50.

### Footprint analysis

To determine the PROX1 footprint, 11 bp-wide motif peaks (CIS-BP ID: M03445_2.00, mapped on mm10 or hg38) were intersected with CUT&RUN consensus peaks, respectively, using bedtools intersect -wa. This generated peaks present in the CUT&RUN consensus peak set that contain motifs and are centred on the motif. DiffTF (v1.8)^106^ was then run using ATAC-seq bam files and the motif-centered peak set^106^.

### Single-cell RNA-seq multiplexing and library generation

For single-cell RNA-sequencing, reprogrammed cells were dissociated into single cells after 2 PBS washes by Accutase digestion. Cells were lifted using cell scrapers, then suspensions were then passed through 70 μm filters into 9 mL of pre-warmed MEF medium. Cells were pelleted (800 rpm, 5 min) and resuspended in 1 mL HBSS + 0.04% BSA to > 3.5 million cells/mL, then fixed through the addition of 4× volume of ice-cold methanol to a final concentration of 80% methanol. Cells were then stored at -20° C. Barcoding oligonucleotides for multiplexing were designed based upon the ClickTag scheme^107^ and ordered with a 5’ amine group label (**Supplementary Table 12**). A methyltetrazine group was conjugated to the oligos via the 5’-amine group. Membrane proteins on the fixed cells were conjugated to an amine-*trans*-cyclooctene group. Subsequent incubation of oligos and cells according to the ClickTag protocol^107^ allowed chemical labelling of cells. ~10,000 cells were multiplexed and loaded per GEM well in the Chromium Controller (10x Genomics), and the Chromium Single Cell 3’ v2 reagent kit was used according to the manufacturer’s instructions for gene expression library generation. We performed modifications at the cDNA and library preparation steps as suggested in the ClickTag protocol to generate barcoded oligonucleotide libraries in parallel. Gene expression and barcode libraries were then diluted to equimolar amounts, pooled at a 9 to 1 ratio, and 26 + 98 bp paired-end sequencing was performed using NovaSeq 6000 platform (Illumina).

### Single-cell RNA-seq data processing

Reprogramming single-cell RNA-seq data generated in this study was analysed using 10x Genomics Cell Ranger (v4.0.0) and Seurat (v4.3)^108^. Cells containing fewer than 1,000 features, containing fewer than 2,000 reads, or with mitochondrial genes comprising over 20% of genes, were discarded. Cell doublets were removed using Scrublet^109^ with a threshold of 0.35. Cells were demultiplexed based on their hashtag oligos by running Seurat HTODemux() recursively. Expression values of cell cycle genes were regressed out using vars.to.regress in ScaleData() to reduce heterogeneity caused by cycling cells. Cells were projected into 2-dimensional space using the UMAP algorithm. Cells were clustered using the Leiden clustering algorithm with 40 dimensions and a resolution parameter of 0.35.

### Regulon analysis

The PROX1 regulon was defined as genes that exhibited downregulation in PROX1 repressor fusion or upregulation in activator fusion, and contained a PROX1 CUT&RUN peak and motif within 1 kb of the TSS. For all other transcription factors Dorothea (v1.7.2, all confidence levels)^110^ was used to build their target gene regulon. Genes in each regulon can be found in **Supplementary Table 8**. We constructed a sub-gene-regulatory network containing the PROX1 regulon and the regulons of transcription factors that are directly regulated by PROX1. GRaNPA was used to determine the most important transcription factors based on this sub-gene-regulatory network and differential expression analysis between *Prox1* overexpression and control^50^.

### Activity score quantification for 4in1 TFs and Prox1

4in1 transcription factors activity was inferred based on the aggregate expression of all genes within the 4in1 regulon from Dorothea, calculated using Seurat addModuleScore(). PROX1 activity was calculated by taking the inverse of the aggregate expression, calculated with addModuleScore(), of a subset of the PROX1 regulon that consists of 79 high-confidence PROX1 repressed target genes (**Supplementary Table 8**). These contained a PROX1 CUT&RUN peak and motif within 1 kb of the TSS and their promoters were differentially closed at day 2 following PROX1 overexpression based on ATAC-seq log2FC threshold (GFP vs *Prox1*) of > 1.

### Filtering cells without transduction

We used PROX1 activity as a basis for the removal of cells from the Prox1 condition which were not successfully transduced with *Prox1* overexpression lentivirus. For each Prox1-labelled cell, we determined the proportion of GFP-labelled cells with a lower PROX1 activity than that cell (GFP_proportion_), and the proportion of Prox1-labelled cells with a higher PROX1 activity than that cell (Prox1_proportion_). We then excluded cells in which GFP_proportion_ was larger than Prox1_proportion_. A similar process was employed using 4in1- and MEF-labelled cells to remove cells from the dataset that were not transduced with 4in1 overexpression lentivirus.

### Signature gene analysis

For analysis of cell type gene signatures, marker genes from the Panglao database^111^ were used as input with Seurat addModuleScore(). The Pearson correlation coefficient between cell type geneset scores and 4in1 or PROX1 activity across all cells was used as the correlation score between cell identity and transcription factor activity.

### Reanalysis of Myc-driven liver tumour and DDC-liver injury scRNA-seq data

Published Myc-driven liver tumour single-cell RNA-seq^37^ data was re-analysed using Seurat along with the provided metadata. After data normalisation with SCTransform, we performed dimensional reduction using PCA and UMAP, using the top 30 dimensions. Clustering was then carried out using the Leiden clustering algorithm with a resolution of 0.2. To ascertain cell identity, we calculated scores based on the Panglao dataset using Seurat module scores. For our specific study objectives, we narrowed our focus to wild type samples classified as “healthy” and those from day 28 upon Myc-overexpression. Published DDC-liver injury single-cell RNA-seq data^39^ was re-analyzed using Seurat in conjunction with provided metadata, focusing on cells generated using the 10x platform and samples from DDC-injected subjects. After data normalisation with SCTransform, dimensional reduction was carried out using UMAP and PCA based on 50 dimensions. We used predefined cell type annotations by the main authors and calculated cell identity using Seurat module scores with reference to the Panglao dataset. Pseudotime trajectory was defined using Monocle3. Subsequently, we analysed *Prox1* expression and hepatocyte identity scores across this trajectory.

### Statistics

Data are presented as mean ± SD. No statistical methods were used to predetermine the sample size. The exact number of technical and biological replicates as well as the applied statistical tests are indicated in the figure or in the figure legend. The biological replicates for all sequencing experiments are listed in **Supplementary Table 13**.

### Survival analysis

For all survival analyses, including patients and mouse models, survival durations were plotted as Kaplan-Meier curves, and a log rank test used to analyse the statistical significance of differences in survival outcomes. For HCC patient survival impact analysis in **Fig. 1e**, -log10(p-values) from the log rank test are shown, with values positive if survival was improved with higher candidate expression, and negative if survival was poorer.

### Plasmid constructs

DNA constructs were generated by DNA synthesis (Sigma) or PCR amplification of cDNA with Q5 polymerase followed by ligation into restriction-digested vectors using indicated enzymes and T4 DNA ligase (NEB). All constructs and primers generated in this study can be found in **Supplementary Tables 10 and 12**, respectively.

### qPCR primers

DNA oligonucleotide primers for quantitative PCR were ordered from Sigma. All primers used in this study are described in **Supplementary Table 12**.

### Antibodies

All primary antibodies used in this study can be found in **Supplementary Table 11**. Secondary Alexa-conjugated antibodies for immunofluorescence were used at 1:2,000 (Invitrogen) and secondary IRDye-conjugated antibodies for Western blot were used at 1:10,000 (LI-COR).

## Acknowledgments

The authors thank L. Butthof, A. Seretny, F. Müller, S. Prokosch, U. Rothermel, J. Hetzer, T. Machauer, J. Hu, and L. V. C. Marques for technical support, and C. Arnold for analysis pipelines and advice. We also thank the DKFZ core facilities, specifically the scOpenLab (P. Mallm), Sequencing Open Lab (N. Glaser), and Genomics and Proteomics Core Facility (A. Schulz), as well as the Nikon Imaging Centre, University of Heidelberg (C. Ackermann), for providing excellent service. We thank Duncan Odom and members of the Mall, Zaugg, Heikenwälder, and Tscharhaganeh labs for discussions.

## Author contributions

Conceptualization: M.M., J.B.Z.; *In Vitro* Experimentation: B.L., J.A.S., B.G.R, L.D., E.P., K.K., M.R.; Software: A.K., I.L.I.; Data Analysis: A.K., B.L., I.L.I., B.G.R, M.R., E.P., I.B., M.S., D.H.; Visualisation: B.L., A.K., B.G.R; *in vivo* Experimentation: K.V., L.B., B.G.R., T.K.; Histology: D.H., B.G.R., B.L., M.R., S.G., J.B.A., D.F.T., H.W.; Resources: J.B., D.G., S.S., T.M., M.B., M.H., D.F.T., H.W.; Funding Acquisition: M.M., J.B.Z.; Supervision: M.M., J.B.Z.; Writing – original draft: M.M., B.L., A.K., J.B.Z..

## Correspondence

Correspondence and requests for materials should be addressed to M.M. or J.B.Z.

## Data availability

All data are present in the manuscript and the supplementary materials. Raw mass spectrometry data have been deposited to the ProteomeXchange Consortium via the PRIDE partner repository. Raw next-generation sequencing data can be found on GEO.

## Code availability

All code used in this study is available from the authors upon request.

## Supplementary Information

Supplementary Information is available for this paper.

Extended Data Figs. 1 – 15

Supplementary Table 1 – 13

References (*70 – 111*)

## Supplementary Figures

**Extended Data Fig. 1:**
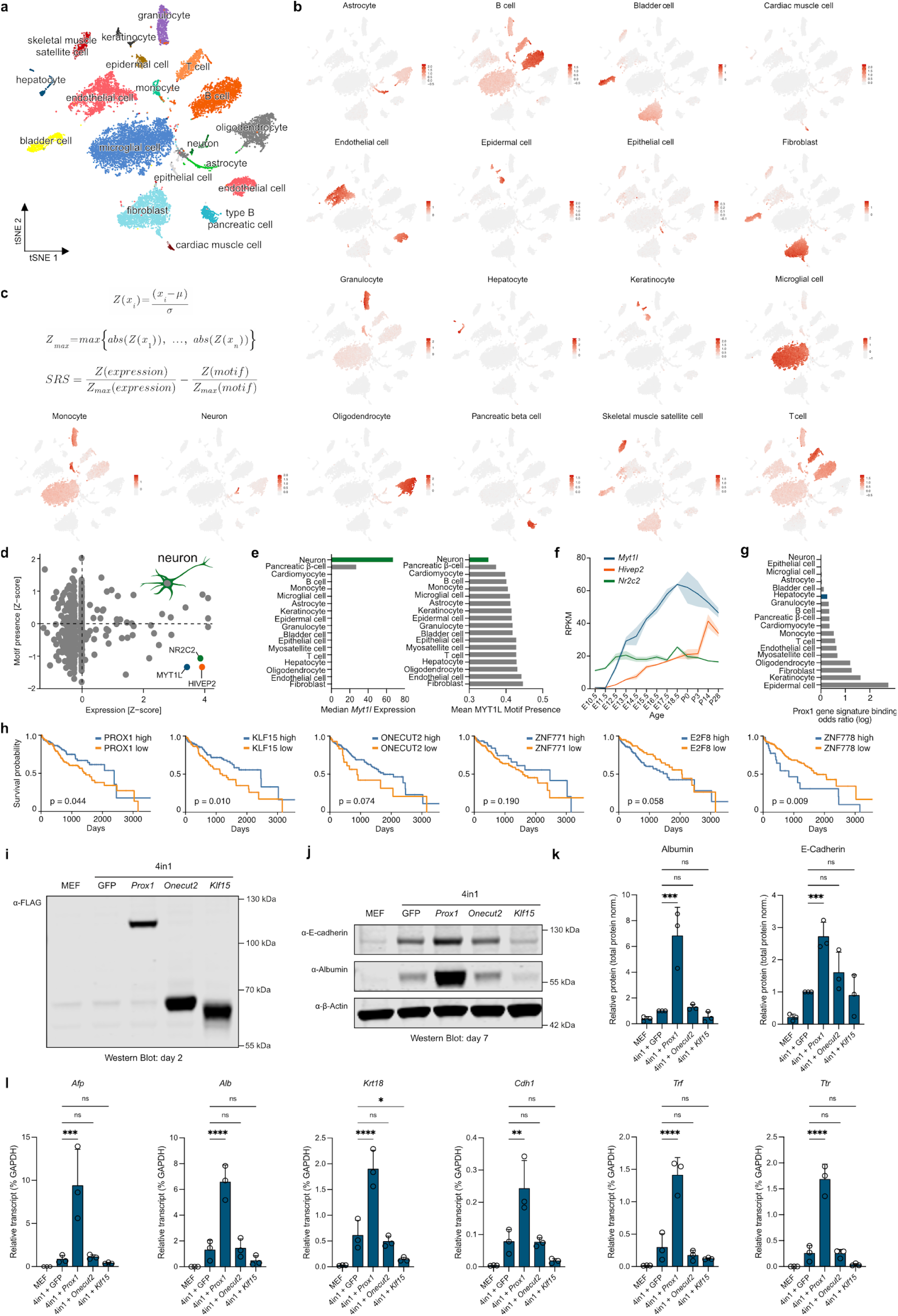
*In silico* and reprogramming screen identifies safeguard repressors. a, Single cell UMAP of 18 cell types annotated by the Tabula Muris consortium. b, Cell type-specific gene signatures used in this study are displayed across all cells in (a). c, Equations used to calculate a safeguard repressor score for each transcription factor (see **Methods** for details). d, Expression and motif presence analysis of 1,296 transcription factors highlights three safeguard repressor candidates in neurons, including MYT1L. e, *Myt1l* expression in 18 cell types from Tabula Muris and number of MYT1L motifs in promoters of cell type marker genes. f, Developmental expression levels of the top three neuronal safeguard repressor candidates in the mouse brain^18^. g, Odds ratio of PROX1 CUT&RUN peaks from mouse liver at promoters of cell type marker genes. h, Kaplan-Meier survival curves for HCC patients in TCGA, segmented by high (blue) or low (orange) expression of each of the candidate hepatocyte safeguard repressors. Log rank p-values are shown. i, Detection of FLAG-tagged hepatocyte safeguard repressor candidates upon overexpression in MEFs at day two of hepatocyte reprogramming using anti-FLAG Western blot. j, Western blot analysis of Albumin and E-cadherin protein levels at day 7 of hepatocyte reprogramming with indicated hepatocyte safeguard repressor candidates. k, Quantification of Albumin and E-cadherin protein expression in (j), normalised to total protein expression. l, Gene expression analysis of indicated hepatocyte markers in cells treated as in (j) using qRT-PCR. Bar graphs show mean values from three biological replicates, error bars = SD, Dunnett’s test, * p-adj < 0.05, ** p-adj < 0.01, *** p-adj < 0.001, **** p-adj < 0.0001.

**Extended Data Fig. 2:**
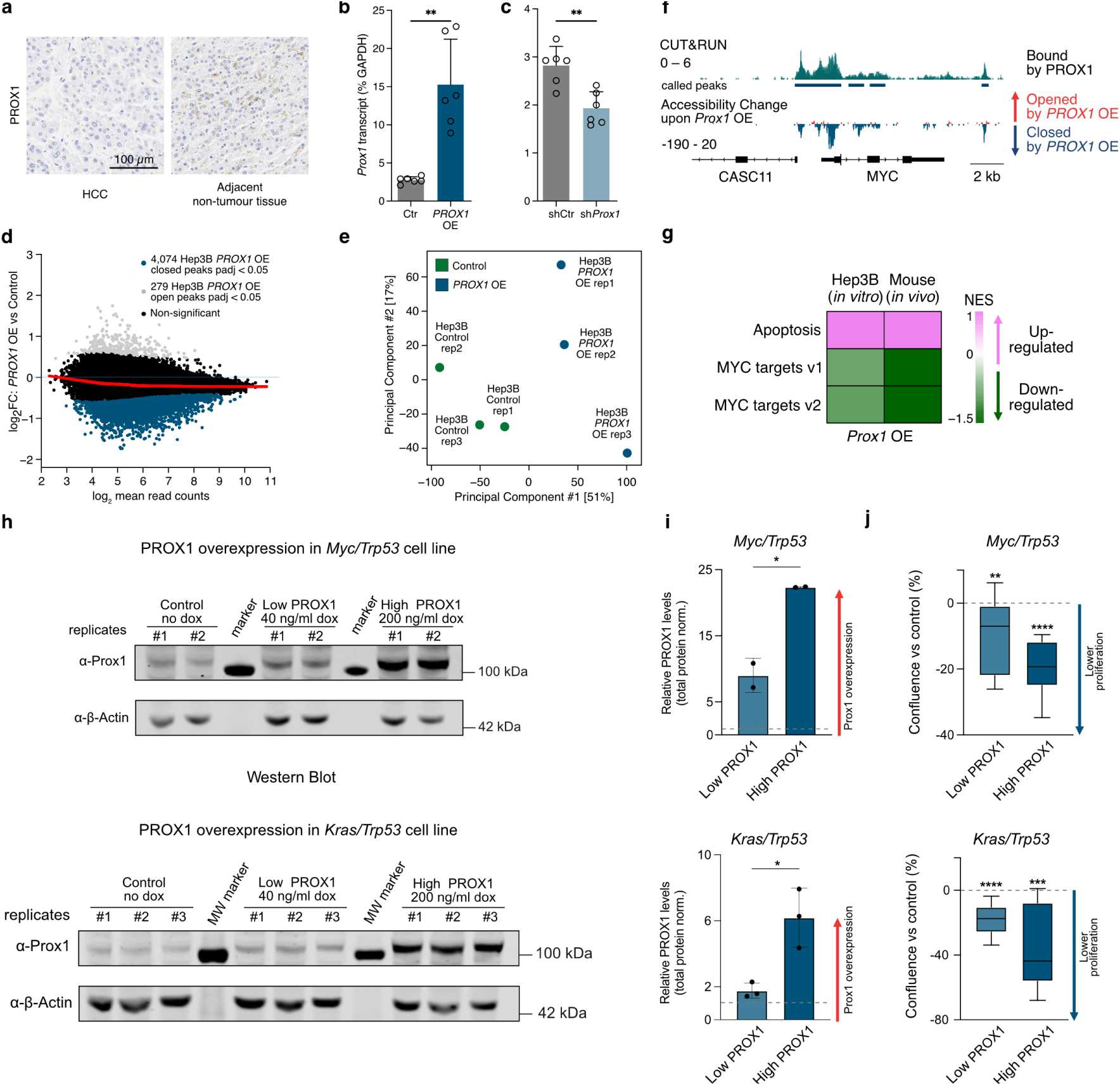
PROX1 is low in HCC tumours and overexpression induces chromatin closure and dose-dependent growth delays in mouse HCC cell lines. a, Representative PROX1 histological micrographs of livers from HCC patients with areas depicting tumour and non-tumour tissue, showing no PROX1 expression in the tumours. b, *PROX1* overexpression following stable integration into Hep3B cell lines and doxycycline treatment for two days (PROX1 OE) compared to control (Ctr) determined by qRT-PCR (n=6). c, Inducible shRNA-mediated *PROX1* knockdown upon doxycycline treatment for two days in Hep3B cell lines (shPROX1) compared to control (shCtr) determined by qRT-PCR (n=6). d, MA-plot of differential accessible regions (DAR) following two days of PROX1 overexpression in Hep3B cells, based on ATAC-seq. e, PCA of DARs in (d) labelled by condition and replicate. f, IGV tracks of PROX1 binding by CUT&RUN and chromatin accessibility at the MYC locus based on ATAC-seq in Hep3B cells two days after PROX1 overexpression displayed as accessibility change compared to control. g, GSEA normalised enrichment scores (NES) for MYC targets and Apoptosis following 7 days of PROX1 overexpression in Hep3B cells, or two days after inducible-*Prox1* overexpression in *Myc/Trp53* HCC mouse model (day 16 harvest). h, Doxycycline dose-dependent *Prox1* overexpression in primary mouse tumour-derived cell lines transformed with *Trp53* knockout together with overexpression of *Myc* (*Myc/Trp53*) (n=2) or *Kras*(G12D) (*Kras/Trp53*) (n=3) determined by Western blot following three days of doxycycline treatment. i, Quantification of PROX1 protein levels of cells treated as in (h). j, Confluency of cells treated as in (h) normalised to uninduced controls. Bar and boxplot graphs show mean values from indicated biological replicates, error bars = SD, two-sided t-test (b,c), unpaired t test (i) or one-sample t-test assuming 0 as mean for (j), * p-adj < 0.05, ** p-adj < 0.01, *** p-adj < 0.001, **** p-adj < 0.0001.

**Extended Data Fig. 3:**
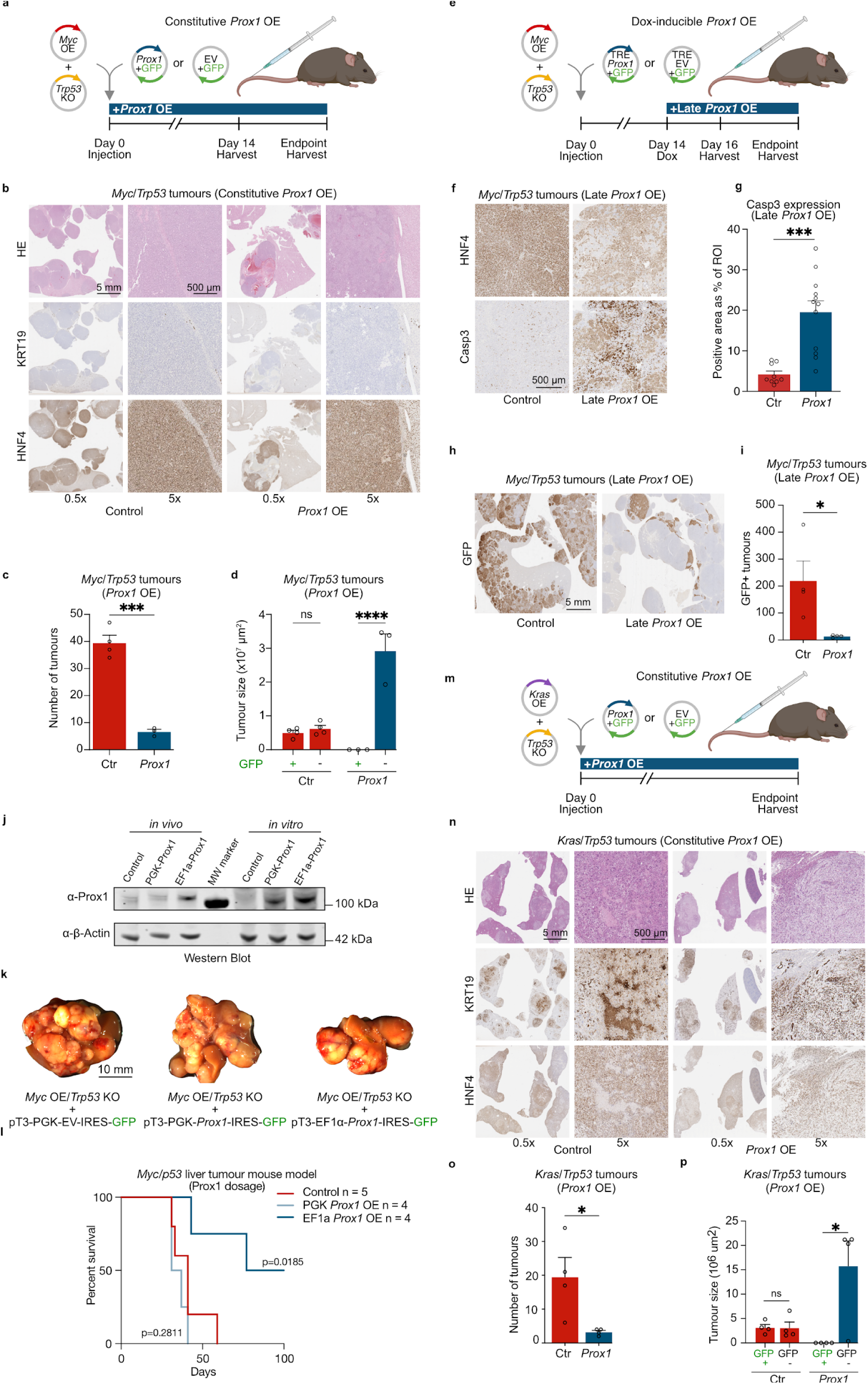
PROX1 prevents liver cancer induction and progression in mice. a, Strategy to induce HCC-like tumours by stable *Myc* overexpression and *Trp53* knockout using HDTVI together with constitutive *Prox1* overexpression compared to GFP controls. b, Representative hematoxylin and eosin (HE), HNF4, and KRT19 histological micrographs of livers treated as in (a) and **Fig. 2h** at both 0.5x and 5x magnification, showing high HNF4 and low KRT19 expression. c, Number of tumours upon treatment as in (a) following overexpression (OE) of *Prox1*-IRES-GFP (n=3) vs GFP control (n=4). d, Size of tumours (cross-sectional area) in (c) with or without GFP expression indicating transgene expression. e, Schematic of doxycycline-inducible Late *Prox1* OE at day 14 following HDTVI-tumour induction as in (a). f, Representative mouse livers treated as in (e) following histological staining for HNF4 and CASP3 at day 16. g, Quantification of CASP3 protein levels in GFP+ tumours in (f) shown as percentage, following Late *Prox1* overexpression (OE) (n=3) vs GFP control (n=4). h, Representative GFP staining in livers treated as in (e) at the endpoint. i, Quantification of GFP+ tumour nodule numbers shown in (h). j, Western blot of PROX1 protein levels following PGK- and Ef1a-promoter driven overexpression *in vivo* following HDTVI and *in vitro* following transfection in respective *Myc*/*Trp53* models compared to controls. k, Representative mouse livers following HDTVI to induce *Myc* overexpression and *Trp53* knockout together with *Prox1* overexpression (OE) using PGK- or Ef1a-promoters (n=4) compared to PGK-GFP controls (n=5). l, Overall survival of mice treated as in (k). Log rank test. m, Strategy to induce liver tumours with complex identity following HDTVI-mediated *KrasG12D* overexpression and *Trp53* knockout together with constitutive *Prox1* overexpression compared to controls. n, Representative histology sections of livers treated as in (m) and **Fig. 2k** stained with HE, HNF4, and KRT19 at both 0.5x and 5x magnification, showing high KRT19 and absence of nuclear HNF4 signal in this tumour model. o, Number of tumours upon treatment as in (m) following overexpression (OE) of *Prox1*-IRES-GFP (n=4) vs GFP control (n=4). p, Size of tumours (cross-sectional area) in (o) with or without GFP expression indicating transgene expression. Bar graphs show mean values from indicated biological replicates, error bars = SD, unpaired t-test, * p-adj < 0.05, ** p-adj < 0.01, *** p-adj < 0.001, **** p-adj < 0.0001.

**Extended Data Fig. 4:**
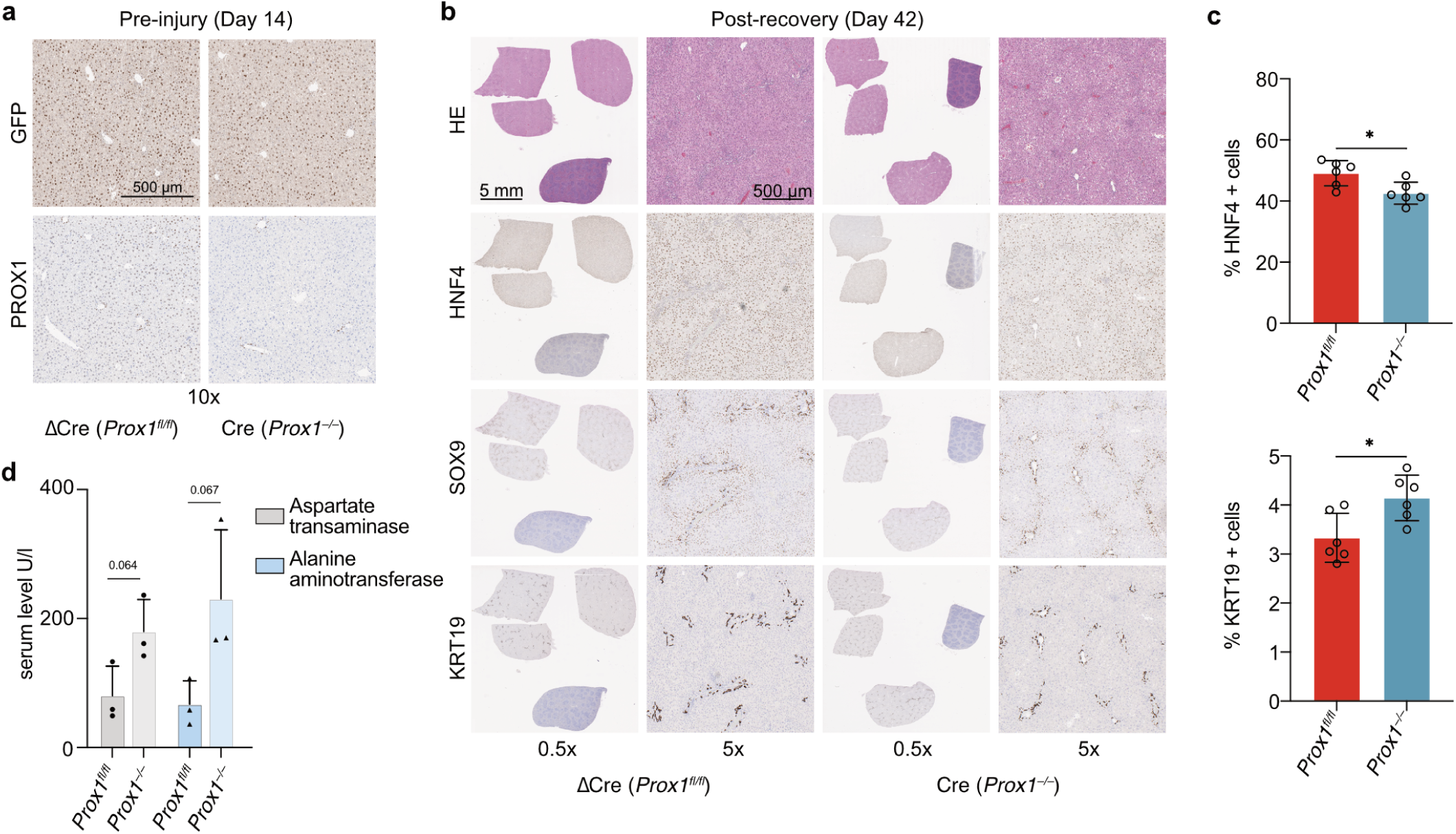
Impaired regeneration upon DDC-liver injury in *Prox1* deleted mice. a, Representative histological staining for PROX1 and GFP of mouse livers two weeks following AAV GFP-cre-mediated *Prox1* deletion in conditional *Prox1* knockout mice (*Prox1^fl^*^/fl^) compared GFP-Δ-cre control mice at day 14 just before injury induction. b, Hematoxylin and eosin (HE), HNF4, SOX9, and KRT19 stainings of liver sections treated as in (a) and following two weeks of DDC-diet and two weeks of recovery with normal diet. c, Percentage of HNF4+ and KRT19+ cells in liver sections from mice treated as in (b) (n=3). d, Serum levels of aspartate transaminase and alanine aminotransferase from mice treated as in (b) (n=3). Bar graphs show mean values from three biological replicates, error bars = SD, unpaired two-sided t-test, * p-adj < 0.05.

**Extended Data Fig. 5:**
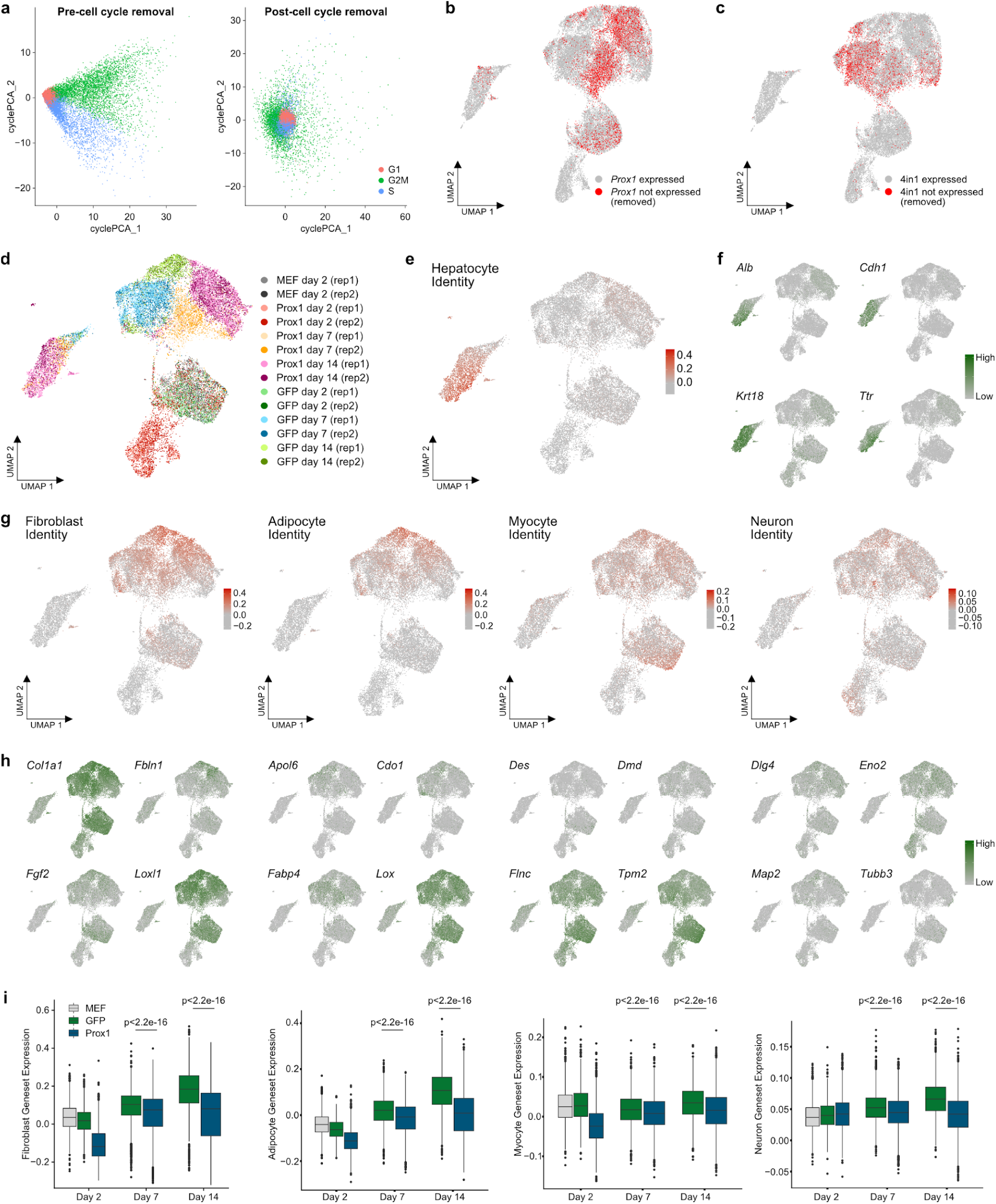
Single cell transcriptome quality control and marker gene expression during hepatocyte reprogramming timecourse with or without Prox1. a, PCA of cells based on cell cycle gene expression before and after adjustment for cell cycle effects using Seurat. b-c, UMAP of cells labelled by expression of *Prox1* (b) and 4in1 (c), inferred from PROX1 and 4in1 activity. d, UMAP of all cells labelled by condition and biological replicate following demultiplexing using barcode oligos. e, Hepatocyte identity score in each cell on UMAP. f, Expression of selected individual hepatocyte marker genes in all cells. g, Fibroblast, adipocyte, myocyte, and neuron identity scores projected onto the UMAP. h, Expression of selected fibroblast, adipocyte, myocyte, and neuronal marker genes in all cells. i, Quantification of fibroblast, adipocyte, myocyte, and neuron identity scores shown as boxplots with p-values (two-tailed t-test) for each time point and treatment.

**Extended Data Fig. 6:**
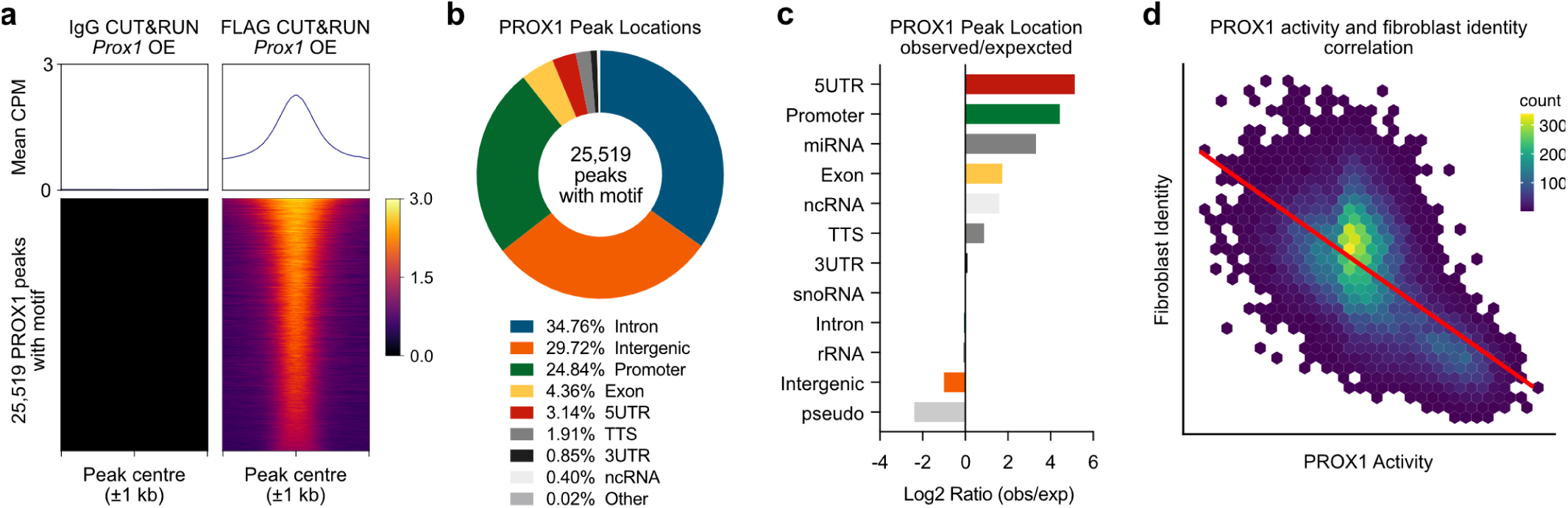
PROX1 chromatin binding sites are enriched at gene promoters. a, PROX1 chromatin binding was determined by CUT&RUN using FLAG antibodies upon overexpression of FLAG-tagged PROX1 in MEFs and compared to GFP overexpression as control identifying 25,519 high-confidence PROX1 binding sites. CUT&RUN using IgG antibodies served as an antibody control and showed no signal at these peaks. b, Proportion of PROX1 binding sites located in indicated genomic regions. c, Log2 ratio of observed PROX1 binding sites to expected binding events in indicated genomic regions highlight enrichment at promoters. d, Negative correlation of fibroblast identity score with PROX1 activity across all single cells during hepatocyte reprogramming time course. p-value < 2.2e-16.

**Extended Data Fig. 7:**
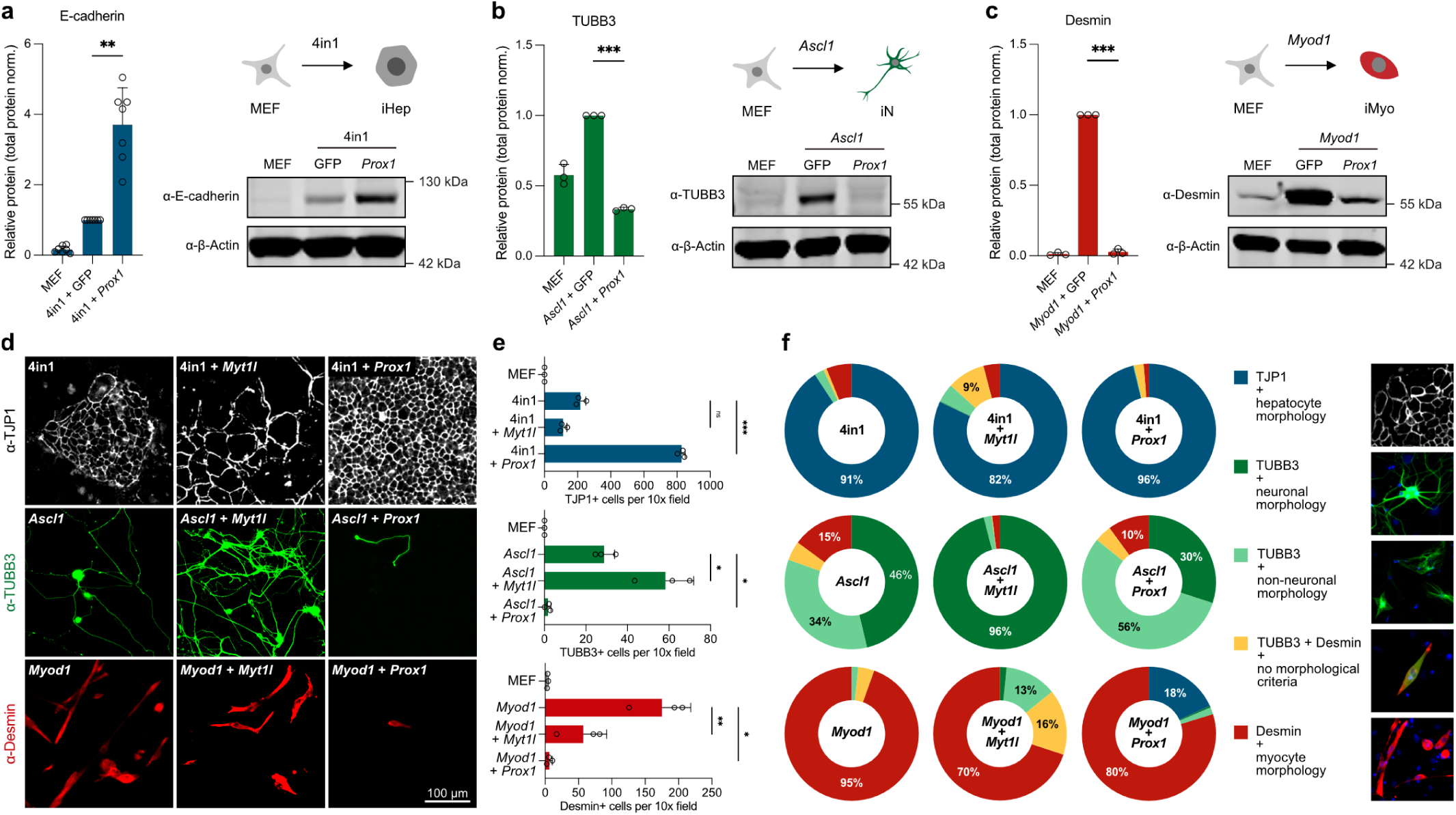
PROX1 inhibits reprogramming to myocyte and neuronal cell fates. a-c, Reprogramming of MEFs to induced hepatocytes, neurons, or myocytes using 4in1, *Ascl1*, or *Myod1* overexpression, respectively. Protein quantification of E-cadherin (hepatocyte), TUBB3 (neuronal), or Desmin (myocyte) marker proteins at day 14 of respective reprogramming protocols with or without *Prox1* overexpression using Western blot analysis. d, Representative immunofluorescence microscopy of cells reprogrammed as in (a-c) with or without *Prox1* or *Myt1l* overexpression in respective reprogramming protocols using indicated antibodies. e, Quantification of successfully reprogrammed cells in (d), based on displayed immunofluorescence markers and morphology. f, Percentage of reprogrammed cells in (d) positive for indicated markers and morphological criteria. Bar graphs show mean values from seven (a) or three (b, c, e) biological replicates, error bars = SD, Dunnett’s test, * p-adj < 0.05. ** p < 0.01, *** p < 0.001.

**Extended Data Fig. 8:**
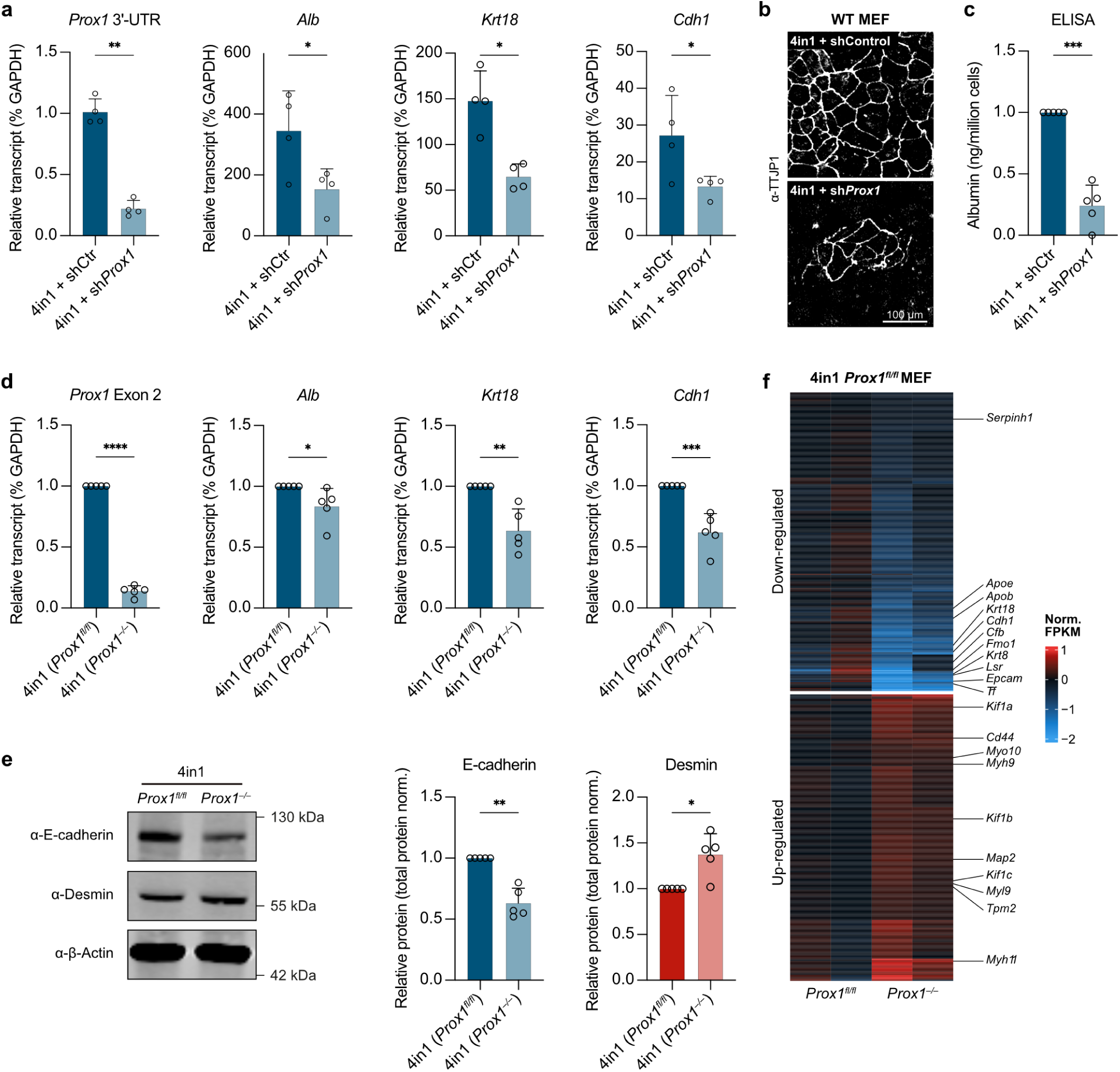
Depletion of *Prox1* reduces hepatocyte reprogramming efficiency and fidelity. a, Expression of *Prox1* and indicated hepatocyte marker genes at day 14 of 4in1-induced hepatocyte reprogramming upon treatment with *Prox1* or control shRNAs based on qRT-PCR. b, Representative TJP1 immunofluorescence of cells in (a) at day 14. c, Quantification of normalised Albumin secretion in cells from (b). d, Expression of *Prox1* and indicated hepatocyte marker genes at day 14 day of hepatocyte reprogramming using *Prox1^fl^*^/fl^ MEFs upon treatment with cre (*Prox1*^−/−^) or Δcre (*Prox1^fl^*^/fl^) as isogenic control based on qRT-PCR. e, E-cadherin (hepatocyte) and Desmin (muscle) protein quantification of cells in (d) based on Western blot analysis. f, Differential gene expression of cells in (d) based on RNA-seq (n=2) highlights downregulation of hepatocyte-specific genes, such as *Krt18* and *Trf*, and upregulation of alternate fate-specific genes, such as *Myh9* (muscle) and *Map2* (neuron), upon *Prox1* deletion. Bar graphs show mean values from four (a) or five (c-e) biological replicates, error bars = SD, two-tailed t-test, * p-adj < 0.05, ** p-adj < 0.01, *** p-adj < 0.001, **** p-adj < 0.0001.

**Extended Data Fig. 9:**
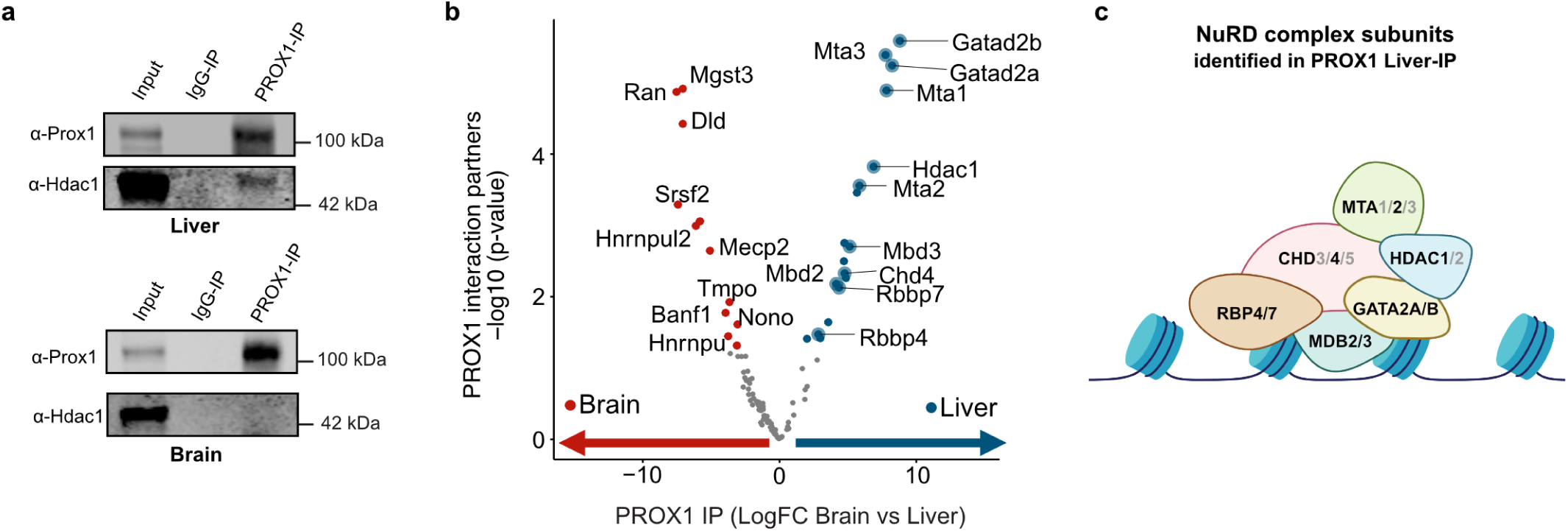
PROX1 interacts with the repressive NuRD complex in liver but not in hippocampus. a, Immunoprecipitation (IP) of PROX1 from primary mouse liver and hippocampus compared to IgG control followed by Western blot using indicated antibodies. b, Mass spectrometric identification and analysis of differential PROX1 interaction partners between hippocampus (red, n=4) and liver (blue, n=4) (see **Methods**). c, Cartoon of the repressive NuRD complex highlighting liver-specific PROX1 interaction partners.

**Extended Data Fig. 10:**
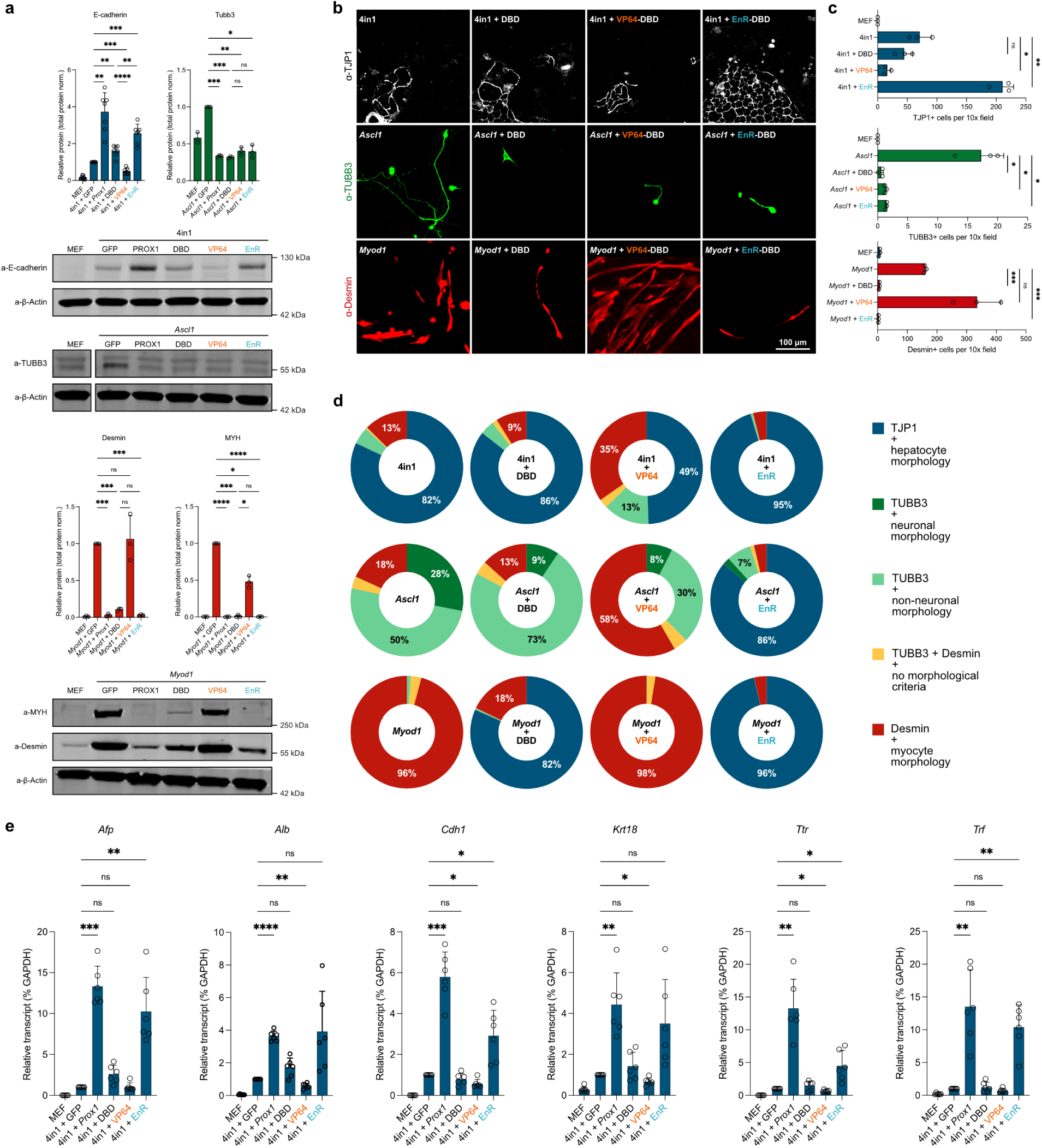
PROX1 target gene repression mimics PROX1 function while target activation has the inverse effect. a, Reprogramming of MEFs to induced hepatocytes, neurons, or myocytes using *MyoD1*, *Ascl1*, or 4in1 overexpression, respectively. PROX1 target gene activation was induced by fusion of PROX1 DNA-binding domain (DBD) to the VP64 activator, while silencing was triggered by EnR repressor fusion. Full-length PROX1 and DBD served as controls. Protein quantification of E-cadherin (hepatocyte, n=7), Tubb3 (neuronal, n=3), or MYH and Desmin (muscle, n=3) marker proteins at day 7 of respective reprogramming experiments using Western blot analysis. b, Representative immunofluorescence microscopy images of cells in (a) at day 14 of reprogramming using indicated antibodies. c, Quantification of successfully reprogrammed cells in (b) based on displayed immunofluorescence markers and morphology. d, Percentage of reprogrammed cells in (b) positive for indicated markers and morphological criteria. e, Expression of indicated hepatocyte marker genes at day 7 of 4in1-induced hepatocyte reprogramming upon overexpression of indicated PROX1 fusion constructs based on qRT-PCR (n=6). Bar graphs show mean values from specified number of biological replicates, error bars = SD, Dunnett’s test, * p-adj < 0.05, ** p-adj < 0.01, *** p-adj < 0.001, **** p-adj < 0.0001.

**Extended Data Fig. 11:**
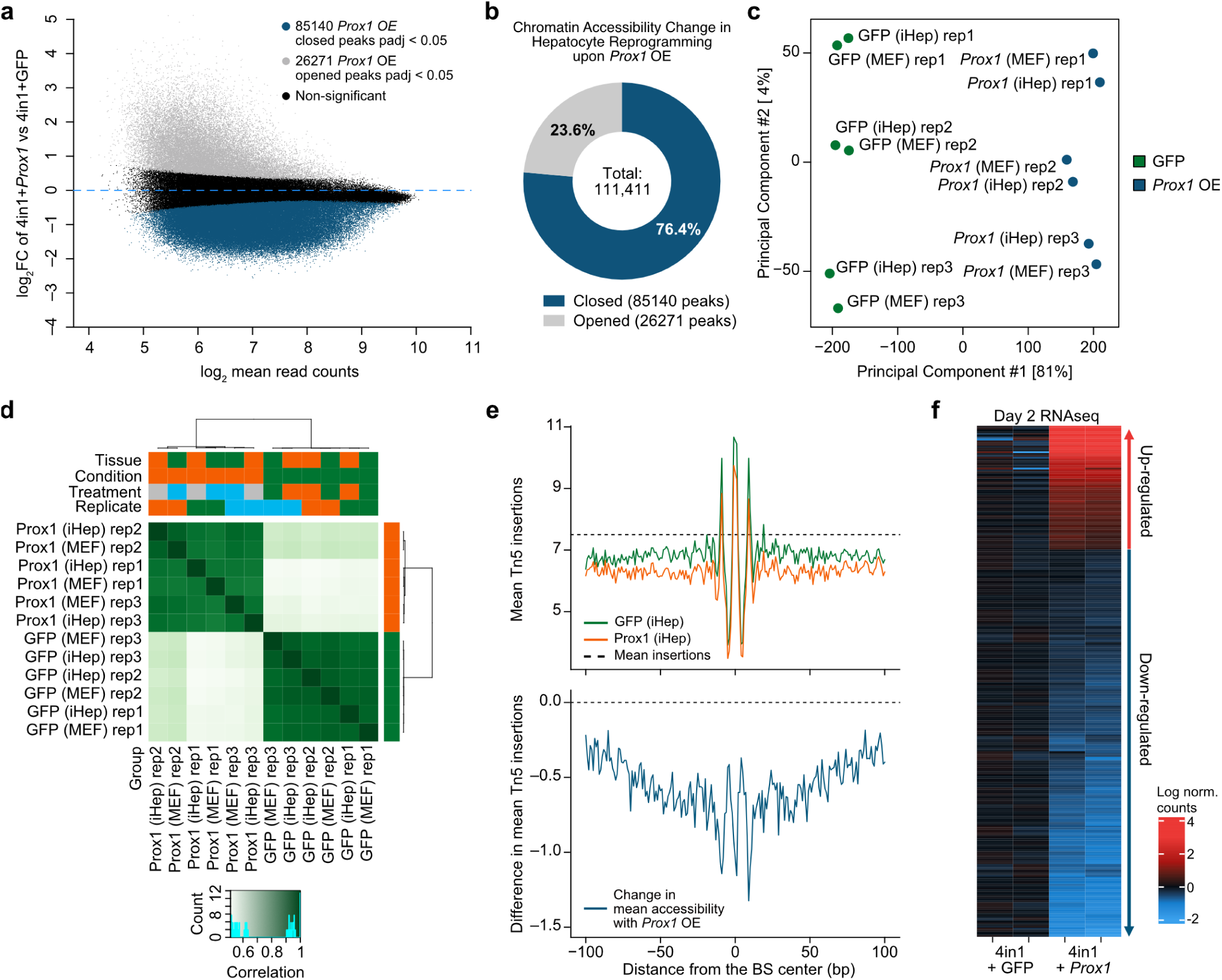
PROX1 predominantly closes bound chromatin and silences associated genes. a, MA-plot of differentially accessible regions (DAR) at day two of 4in1-induced hepatocyte reprogramming with or without *Prox1* overexpression based on ATAC-seq (n=3). b, Percentage of closed and opened regions upon *Prox1* overexpression in (a). c, Correlation of DARs between indicated conditions and replicates in (a) and upon overexpression of *GFP* or *Prox1* in MEFs. d, PCA of DARs in (c) labelled by condition and replicate. e, Top, mean Tn5 transposon adapter insertions at indicated conditions from (a) centred at PROX1 binding sites from CUT&RUN experiments. Bottom, changes in chromatin accessibility at PROX1 binding sites between 4in1 + GFP and 4in1 + Prox1 indicate decreased accessibility upon *Prox1* overexpression. f, Differential gene expression of cells in (a) based on RNA-seq (n=2), for genes with differentially-accessible regions (p-adj < 0.05) within 2 kb of their TSS and an overlapping CUT&RUN peak with a PROX1 binding motif, showing global downregulation of target gene transcription upon *Prox1* overexpression.

**Extended Data Fig. 12:**
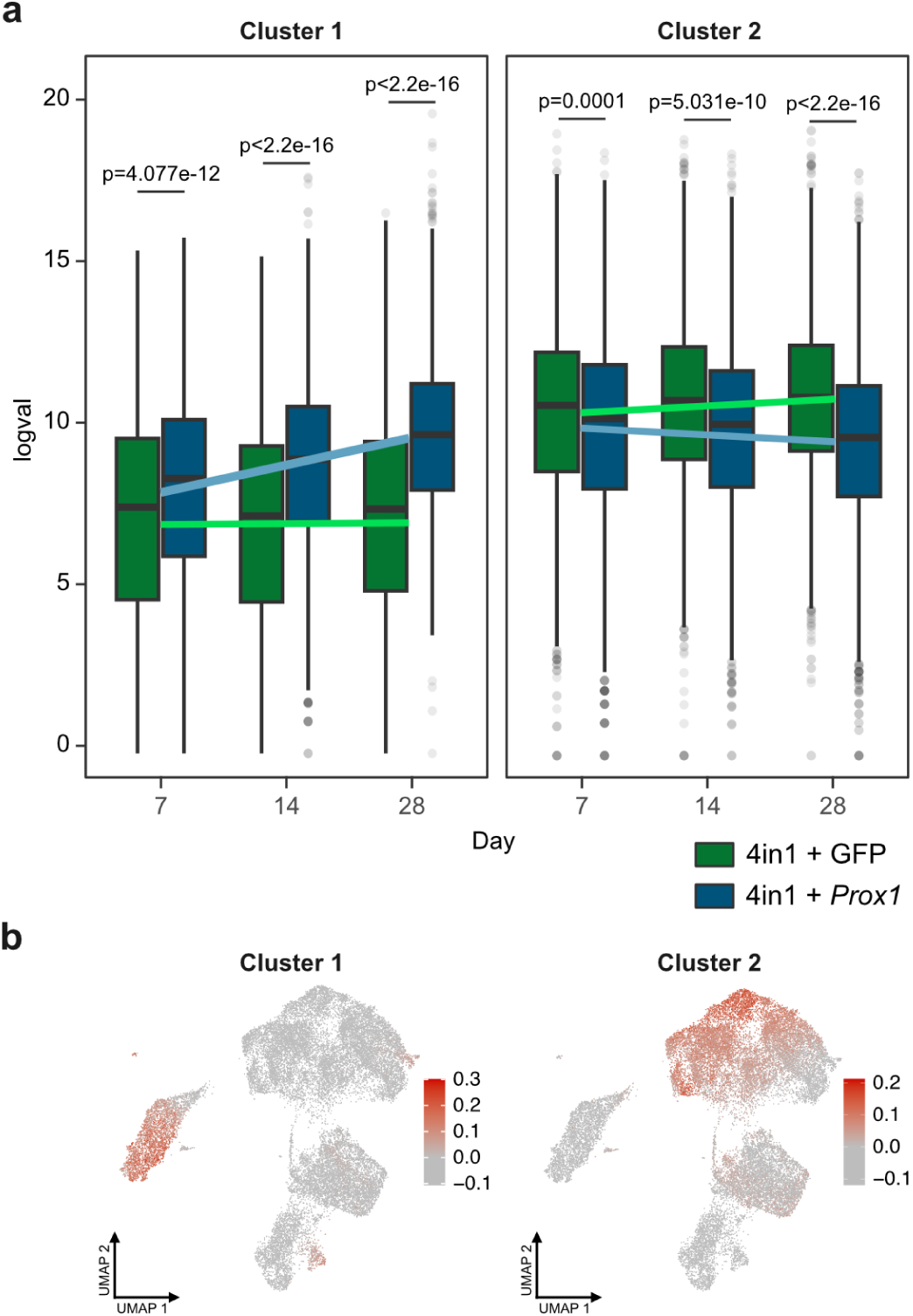
Induction of hepatocyte gene cluster 1 and repression of alternate fate cluster 2 by PROX1 in bulk and single cell gene expression datasets over time. a, Bulk gene expression changes during 4in1-induced hepatocyte reprogramming at indicated time points clustered by differential expression and chromatin accessibility. Median gene expression of differential expressed genes between Prox1 and GFP control are presented as boxplots for both clusters, two-tailed t-test (p-values shown). b, Aggregate expression of genes from bulk gene expression-derived cluster 1 and 2 projected onto UMAP of 4in1-induced reprogramming single-cell transcriptomics data from **Fig. 2b**, highlight overlap of cluster 1 with hepatocyte cluster and cluster 2 with alternate fate cluster.

**Extended Data Fig. 13:**
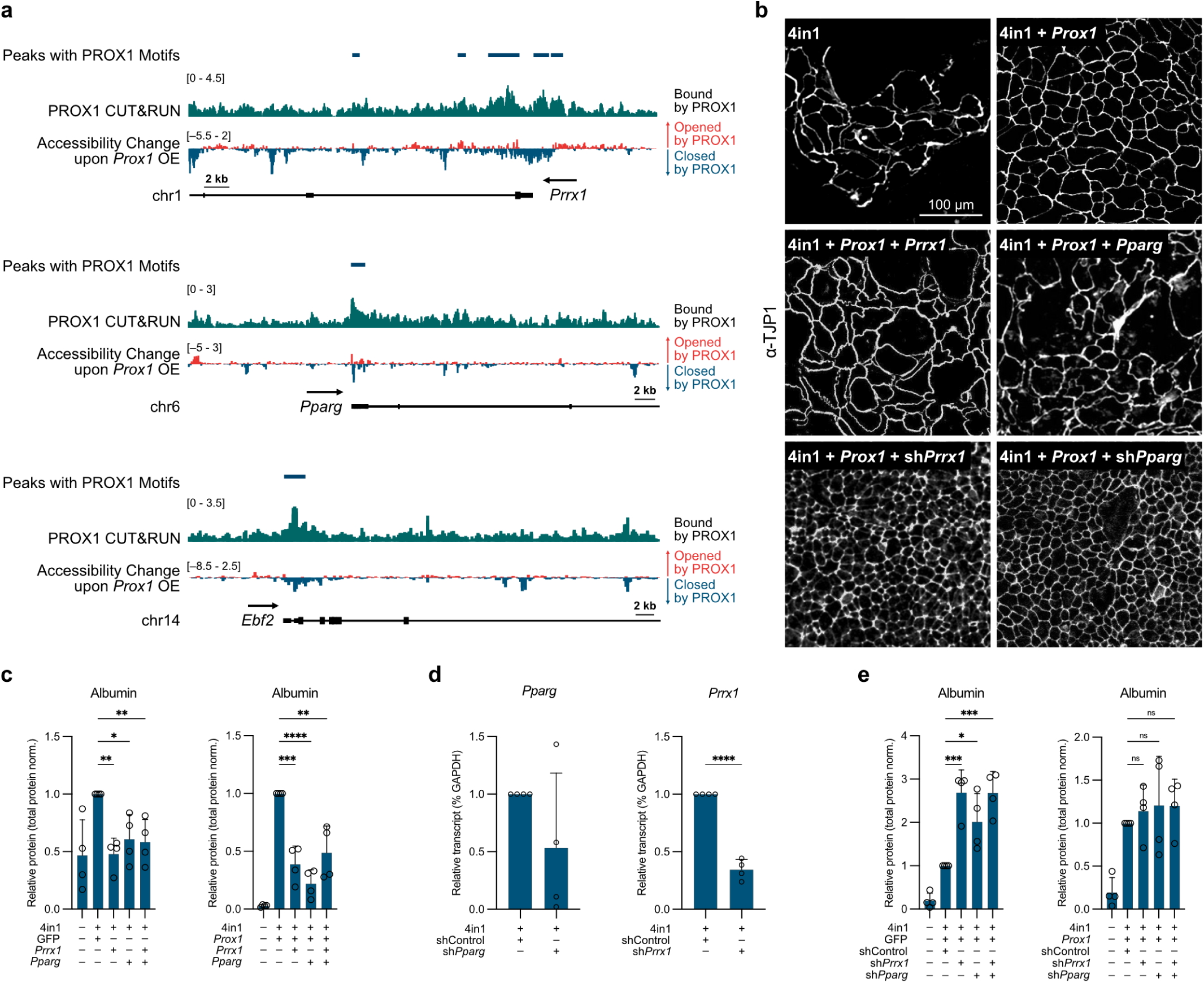
*Prrx1* and *Pparg* are PROX1 target genes and their repression in part explains PROX1 activity during reprogramming. a, Representative IGV tracks of PROX1 chromatin binding based on CUT&RUN (n=3) and chromatin accessibility based on ATAC-seq (n=3) displayed as accessibility change at day two day of 4in1-induced hepatocyte reprogramming with or without *Prox1* at the promoters of *Prrx1* (top), *Pparg* (middle), *Ebf2* (bottom). PROX1 motifs and CUT&RUN peaks are displayed. b, Representative TJP1 immunofluorescence staining at day 14 of hepatocyte reprogramming with indicated combinations of transcription factor overexpression and/or shRNA-mediated knockdown treatments. c, Quantification of Albumin protein levels by Western blot at day 14 of hepatocyte reprogramming with overexpression of *Prrx1* or *Pparg* and co-overexpression of GFP control (left) or *Prox1* (right). d, Expression of *Pparg* and *Prrx1* upon shRNA-knockdown at day 14 of hepatocyte reprogramming determined by qRT-PCR. e, Albumin protein levels determined by quantitative Western blot at day 14 of hepatocyte reprogramming upon knockdown of *Prrx1* or *Pparg* with or without *Prox1* overexpression. Bar graphs show mean values normalised to total protein levels in (c and e), or GAPDH expression in (d), from four biological replicates, error bars = SD, Dunnett’s test (c and e) or two-tailed t-test (d), * p-adj < 0.05, ** p-adj < 0.01, *** p-adj < 0.001, **** p-adj < 0.0001.

**Extended Data Fig. 14:**
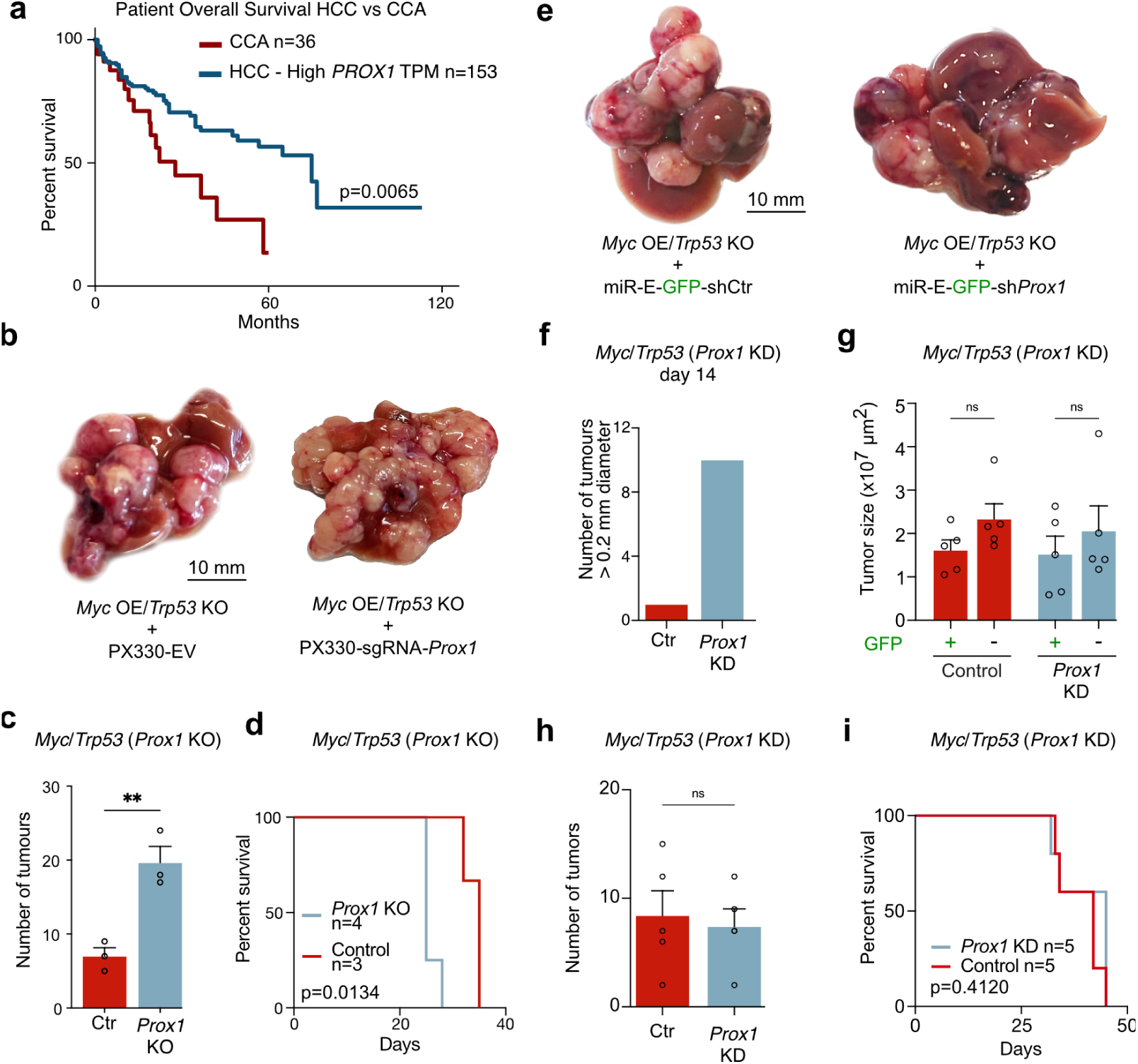
PROX1 loss can enhance HCC liver tumour formation in mice. a, Overall survival of 135 HCC patients with high *PROX1* expression levels (40% cutoff for high-expression cohort) compared to CCA patients^28^. b, Representative photographs of mouse livers at endpoint of HCC tumour modelling induced by *Myc* overexpression and *Trp53* knockout using HDTVI together with PX330-sgRNA-Prox1 knockout compared to respective negative control. c, Number of tumours upon treatment as in (a) following *Prox1* knockout (KO) vs control (Ctr) (n=3). d, Survival curve of HCC mice treated as in (a) comparing *Prox1* KO (n=4) with control (n=3). Log rank test. e, Representative mouse livers at endpoint of HCC tumour modelling as in (a) with EF1a-GFP-shProx1 knockdown compared to respective negative control. Histology sections of corresponding livers are shown in **Fig. 5c**. f, Total number of tumour nodules with diameter > 0.2 mm at day 14 of tumour induction as in (d) upon shRNA-mediated control or *Prox1* knockdown (n=5). g, Tumour size (cross-sectional area) following treatments as in (d) displaying tumours with or without GFP expression (n=5). h, Number of tumours upon treatment as in (a) following *Prox1* knockdown (shProx1) vs control (shCtr) (n=5). i, Survival curve of HCC mice treated as in (d) comparing knockdown of *Prox1* with control (n=5). Log rank test. Bar graphs show mean values from specified biological replicates, error bars = SD, unpaired t-test, * p-adj < 0.05, ** p-adj < 0.01, *** p-adj < 0.001, **** p-adj < 0.0001.

**Extended Data Fig. 15:**
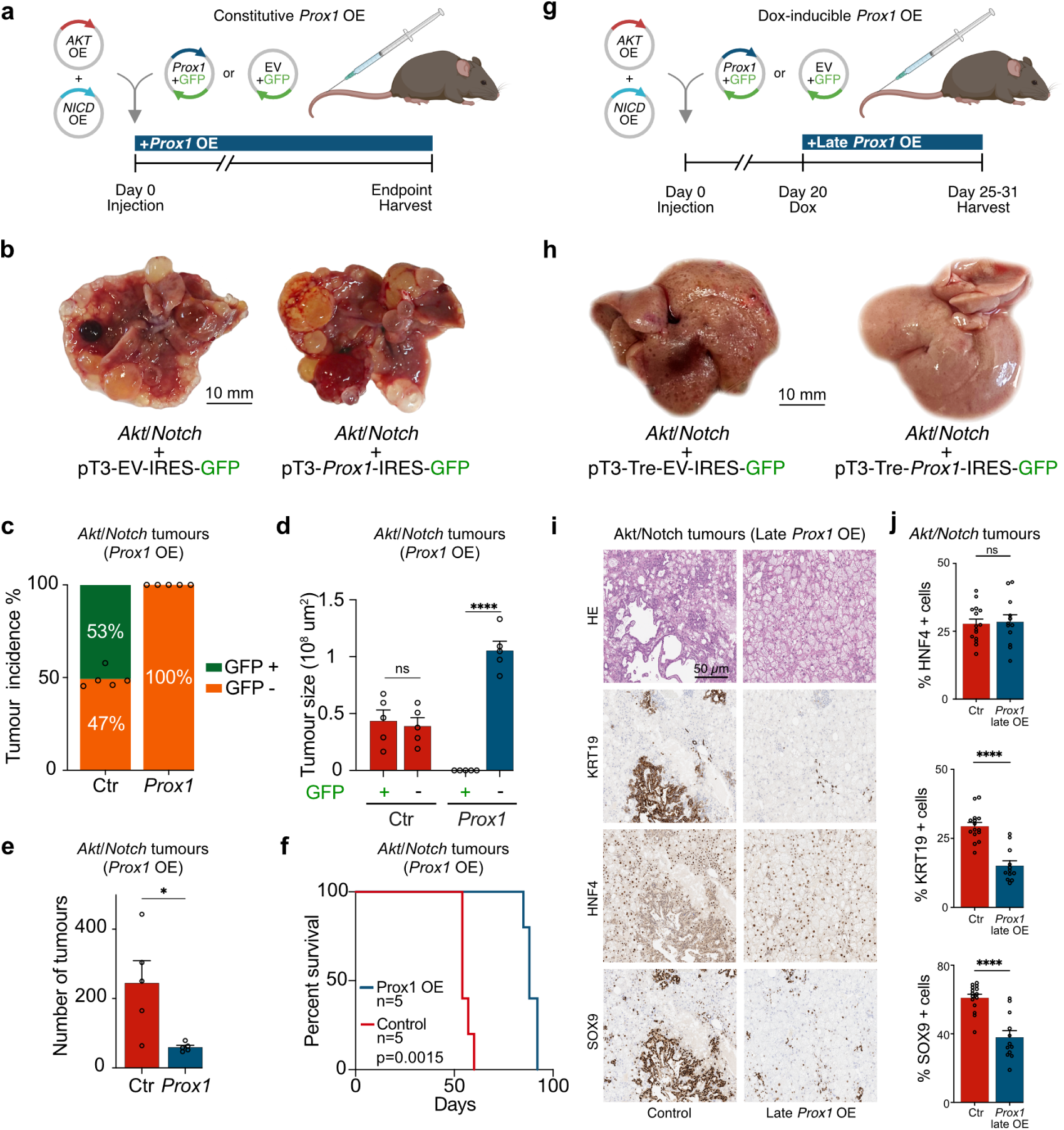
PROX1 can shift liver tumour fate from CCA to HCC in mice. a, Strategy to induce liver tumours with CCA identity following HDTVI-mediated *Akt and Notch1 receptor intracellular domain* (NICD) (*Akt*/*Notch*) overexpression with constitutive *Prox1* overexpression compared to controls. b, Representative mouse livers at endpoint of CCA tumour modelling induced as in (a). Histology sections of corresponding livers are shown in **Fig. 5c**. c, Quantification of GFP + tumours following constitutive overexpression (OE) of *Prox1*-IRES-GFP (n=4) vs GFP control as in (a), shown as percentage (n=5). d, Size of tumours (cross-sectional area) in (c) with or without GFP expression indicating transgene expression. e, Number of tumours upon treatment as in (c). f, Survival curve of CCA mice treated as in (c). Log rank test. g, Schematic of doxycycline-inducible Late *Prox1* OE at day 20 following HDTVI-CCA tumour induction as in (a). h, Representative mouse livers 1-2 weeks following Late *Prox1* OE induced in CCA model as in (g). i, Histology sections of livers treated as in (g) stained with HE, KRT19, HNF4, and SOX9. j, Quantification of KRT19 and SOX9 (CCA-marker) and HNF4 (HCC-marker) positive cells in GFP+ tumours following Late *Prox1* overexpression (OE) (n=4) vs GFP control (n=4) from (g) shown as percentage. Bar graphs show mean values from specified biological replicates, error bars = SD, unpaired t-test, * p-adj < 0.05, ** p-adj < 0.01, *** p-adj < 0.001, **** p-adj < 0.0001.

## Supplementary Tables

Supplementary Table 1: Cell identity gene signatures

Genesets containing marker genes for various cell types, from Tabula Muris and the Panglao database.

Supplementary Table 2: Computational safeguard repressor screen results

Raw and scaled expression and motif density values for all transcription factors and cell types, safeguard repressor scores, lifelong expression predictions, and literature evidence for reprogramming.

Supplementary Table 3: ATAC-seq peaks

Differential chromatin accessibility analysis from: Hep3B cells with *PROX1* overexpression or control at day two; and the hepatocyte reprogramming experiment with *Prox1* overexpression or control, either in MEFs or with 4in1 overexpression at day two.

Supplementary Table 4: PROX1 CUT&RUN

PROX1 CUT&RUN peaks containing PROX1 binding motifs, annotated with closest TSS, from primary mouse liver, Hep3B cells and iHeps upon Prox1 overexpression.

Supplementary Table 5: RNA-seq DEG table in vitro

Differential gene expression analysis from Hep3B cells with *PROX1* overexpression or control

Differential gene expression analysis from the hepatocyte reprogramming experiment with *Prox1^−/−^*^Prox1^ knockout or control MEFs.

Differential gene expression analysis from the hepatocyte reprogramming experiment with *Prox1* DBD fusion proteins. All comparisons were performed with DBD as the baseline control.

Differential gene expression analysis from the hepatocyte reprogramming experiment with *Prox1* overexpression or control, at days 2, 7, 14, and 28.

Supplementary Table 6: Overview of HDTVI experiments Details of mice, including injected plasmids and survival time.

Supplementary Table 7: RNA-seq DEG table in vivo

Differential gene expression analysis from HCC mouse tumours with late *Prox1* overexpression or control. Differential gene expression analysis from *Myc/Trp53* mouse tumours with *Prox1* knockdown or control. Differential gene expression analysis from *Akt/Notch* mouse tumours with late *Prox1* overexpression or control.

Supplementary Table 8: PROX1 target genes

PROX1 regulon gene set and PROX1 activity gene set.

Supplementary Table 9: PROX1 protein-protein interaction partners

Mass spectrometry-based proteomics analysis of PROX1 interaction partners following PROX1 immunoprecipitation from mouse liver and hippocampus.

Supplementary Table 10: Plasmids Plasmids used in this study.

Supplementary Table 11: Antibodies Antibodies used in this study.

Supplementary Table 12: Oligos

qPCR primers, scRNA-seq barcodes, ATAC-seq oligos, and cloning oligos used in this study.

Supplementary Table 13: Description of experimental conditions for all genomics experiments

Experimental conditions, replicates, and associated supplementary table for RNA-seq, scRNA-seq, ATAC-seq, and CUT&RUN experiments.

